# Novel Fas-TNFR chimeras that prevents Fas ligand-mediated kill and signals synergistically to enhance CAR T-cell efficacy

**DOI:** 10.1101/2023.02.22.529492

**Authors:** Callum McKenzie, Mohamed El-Kholy, Farhaan Parekh, Mathew Robson, Katarina Lamb, Christopher Allen, James Sillibourne, Shaun Cordoba, Simon Thomas, Martin Pule

**Author notes:** Correspondence should be addressed to M.P.

## Abstract

The hostile tumour microenvironment limits the efficacy of adoptive cell therapies. Activation of the Fas death receptor initiates apoptosis and disrupting these receptors could be key to increase CAR T-cell efficacy. We screened a library of Fas-TNFR proteins identifying several novel chimeras that not only prevented Fas ligand-mediated kill, but also enhanced CAR T-cell efficacy by signalling synergistically with the CAR. Upon binding Fas ligand, Fas-CD40 activated the NF-κB pathway, inducing greatest proliferation and IFNγ release out of all Fas-TNFRs tested. Fas-CD40 induced profound transcriptional modifications, particularly genes relating to the cell cycle, metabolism, and chemokine signalling. Co-expression of Fas-CD40 with either 4-1BB- or CD28-containing CARs increased *in vitro* efficacy by eliciting maximal CAR T-cell proliferation and cancer target cytotoxicity, and enhanced tumour killing and overall mouse survival *in vivo*. Functional activity of the Fas-TNFRs were dependent on the co-stimulatory domain within the CAR, highlighting crosstalk between signalling pathways. Furthermore, we show that a major source for Fas-TNFR activation derives from CAR T cells themselves via activation-induced Fas ligand upregulation, highlighting a universal role of Fas-TNFRs in augmenting CAR T-cell responses. We have identified Fas-CD40 as the optimal chimera for overcoming Fas ligand-mediated kill and enhancing CAR T-cell efficacy.

## INTRODUCTION

Adoptive transfer of chimeric antigen receptor (CAR) T cells has seen remarkable success in the treatment of relapsed/refractory haematological cancers, however, approximately 60% of patients eventually relapse, partly due to the hostile tumour microenvironment (TME)^1^. Extending these clinical successes to solid tumour indications is more challenging due to an even more complex and immunosuppressive TME^1–3^.

The Fas/Fas ligand (FasL) pathway is a key inhibitory checkpoint contributing to the immunosuppressive TME^4–7^. Fas is a member of the tumour necrosis factor receptor (TNFR) superfamily and comprises one of eight TNFR death receptors^8^. Upon binding FasL, Fas trimerizes allowing for binding of the adaptor protein, Fas-associated death domain (FADD), to the intracellular death domains of Fas via homotypic interactions^9^. Pro-caspase 8 then binds FADD via death effector domains, creating the death-inducing signalling complex, and is then cleaved to activate downstream executioner caspases, initiating apoptosis (Figure 1A).

**Figure 1.**
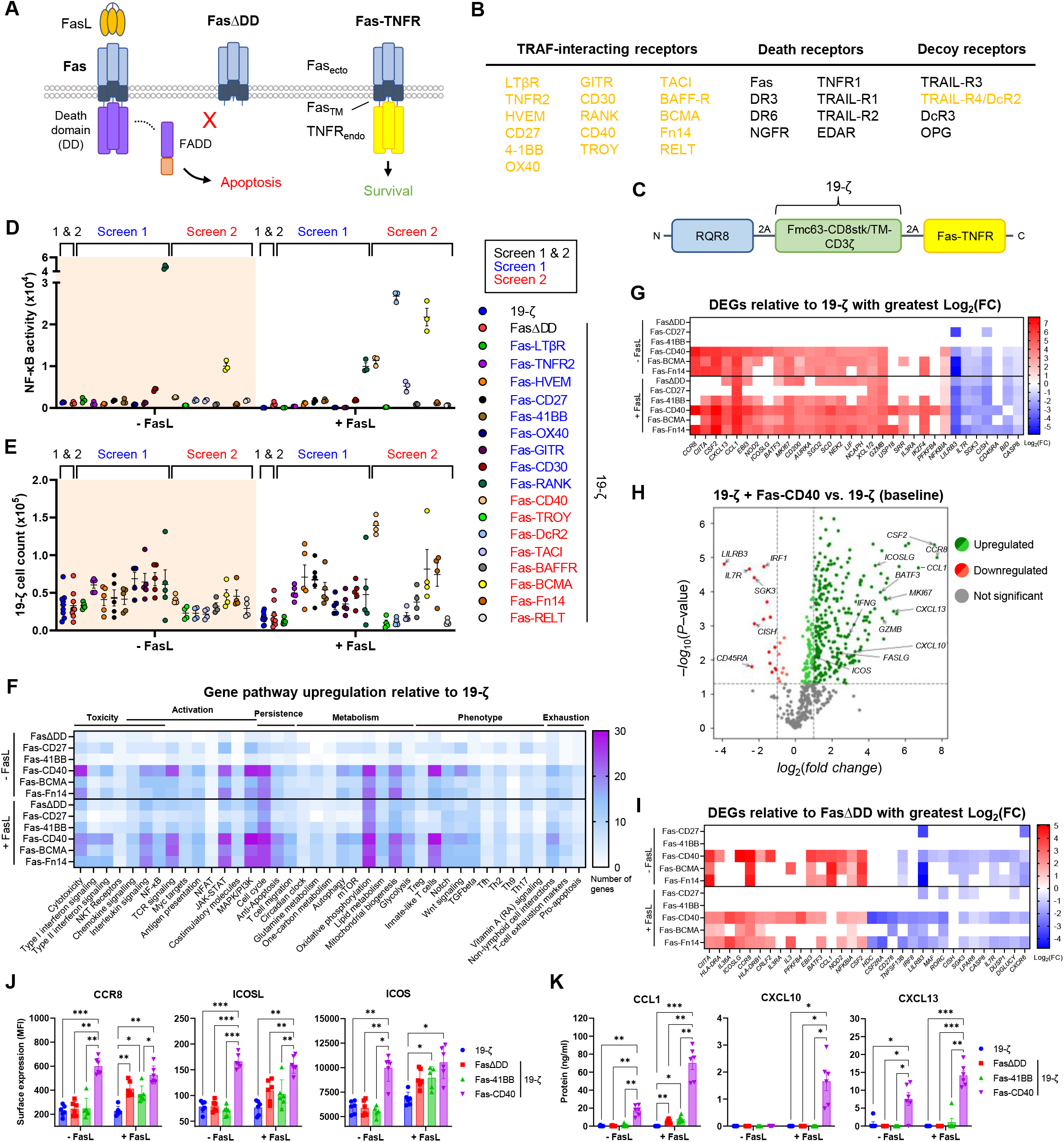
Fas-CD40 activates NF-κB and induces strong proliferation upon FasL binding. (A) Left: Upon binding FasL, Fas trimerisation recruits FADD, initiating apoptosis. Middle: FasΔDD acts as a decoy receptor to FasL by being unable to recruit FADD. Right: Schematic of Fas-TNFR structure; the ectodomain and transmembrane domain of Fas are fused to the endodomains of TNFRs. Upon FasL binding the Fas-TNFR chimera converts the death signal into a survival/growth signal. (B) Members of the TNFR superfamily. Those highlighted in gold were included in the Fas-TNFR screen. (C) Schematic of polycistronic transgene transduced into human T cells. 19-ζ: Fmc63 binder fused to the endodomain of CD3ζ via a CD8 stalk/transmembrane domain. (D) NF-κB reporter Jurkat cells transduced to express either 19-ζ alone or co-express FasΔDD or the Fas-TNFRs were cultured with or without immobilised recombinant FasL (20 μg/ml) overnight, where NF-κB activity was measured. Experiment performed with technical triplicates, error bars are SEM. (E) 5×10^4^ human T cells expressing 19-ζ and FasΔDD or the Fas-TNFRs were cultured with or without immobilised recombinant FasL (20 μg/ml) for five days, at which point cell counts were analysed by flow cytometry. Due to the large number of possible Fas-TNFR chimeras, they were tested over two separate experiments (screen 1 and 2) with the data being compiled onto one graph. Five independent donors were tested in screen 1 and four independent donors were tested in screen 2, error bars are SEM. (F) Human T cells from five independent donors were transduced to express 19-ζ or co-express FasΔDD or the stated Fas-TNFRs and then cultured with or without immobilised recombinant FasL (20 μg/ml) for three days, at which point RNA was extracted and analysed using the nCounter® NanoString® platform with the CAR-T Characterization Panel. 19-ζ cells co-expressing FasΔDD or the Fas-TNFRs were made relative to 19-ζ alone, and the number of significantly (*P* < 0.05) upregulated differentially expressed genes (DEGs) were categorised by pathway involvement. (G) Significant DEGs relative to 19-ζ with greatest Log_2_ fold change (FC) from experiment described in F. (H) Volcano plot from experiment described in F of Fas-CD40-19-ζ cells compared to 19-ζ alone after incubation with immobilised FasL. (I) Significant DEGs relative to FasΔDD-19-ζ with greatest Log_2_(FC) from experiment described in F. (J and K) 19-ζ cells were cultured in the presence or absence of immobilised FasL (20 μg/ml) for five days and then stained for CCR8, ICOSL and ICOS expression by flow cytometry (I), or the cell culture supernatant analysed for CCL1, CXCL10 and CXCL13 secretion (J). Six independent donors tested, error bars are SEM, \**P* < 0.05, \*\**P* < 0.01, \*\*\**P* < 0.001, two-way ANOVA.

T cells constitutively express Fas and are consequently vulnerable to FasL-mediated apoptosis. *FASLG* and FasL are overexpressed by many cancers, either by cancer cells themselves, or by cells constituting the TME, such as regulatory T cells (Tregs), myeloid-derived suppressor cells (MDSCs), cancer-associated fibroblasts (CAFs) and tumour endothelial cells^5,6^. Moreover, T cells upregulate FasL upon activation, inducing fratricide, an effect particularly observed with third generation CARs^10,11^. Therefore, the Fas/FasL checkpoint can limit the efficacy of adoptive T-cell therapy.

Several strategies to overcome FasL in immunotherapy have been explored. Therapeutic monoclonal antibodies which block Fas or FasL effectively prevents FasL-mediated T-cell loss; however FasL-mediated killing of tumour is concomitantly compromized^12–14^. Adoptive immunotherapy with engineered immune cells affords more discrete methods: disruption of Fas expression by small interfering RNAs or CRISPR/Cas9 is effective^15,16^. An alternative strategy is expression of non-functional Fas which outcompetes with native Fas. This latter strategy includes a truncated Fas receptor lacking the death domain (FasΔDD) or a chimeric Fas-41BB protein^5,17–19^. Expression of FasΔDD or Fas-41BB rescues FasL-mediated apoptosis. The Fas-41BB chimera additionally converts the death signal into a pro-survival 4-1BB signal by activating NF-κB and MAP kinase (MAPK) pathways via TNFR-associated factors (TRAFs)^20^.

There are many other members of the TNFR superfamily apart from 4-1BB that provide co-stimulatory signals which due to differential TRAF activation may be qualitatively different. In this paper we performed a functional assessment of Fas-TNFR chimeric proteins in the context of human T cells. We identified several novel Fas-TNFR chimeras that co-stimulates CAR T cells, delivering enhanced target cytotoxicity and CAR T-cell persistence/proliferation compared to FasΔDD and the Fas-41BB chimera. In particular, Fas-CD40 optimally enhanced CAR T-cell efficacy when co-expressed with either 4-1BB- or CD28-containing CARs. Moreover, we demonstrate that a major source of FasL for Fas-TNFR activation derives from T cells themselves, highlighting a universal role of Fas-TNFRs to augment CAR T-cell therapy.

## RESULTS

### Screening of Fas-TNFRs reveals Fas-CD40 as a potent inducer of proliferation upon binding FasL

We first created a set of Fas-TNFR chimeras comprising the ectodomain and transmembrane domain of Fas fused to the endodomains of pro-survival TRAF-interacting TNFRs, as well as the endodomain of the TRAIL decoy receptor, TRAIL-R4/DcR2^21^ (Figures 1A, 1B). The Fas-TNFR chimeras (highlighted in gold in Figure 1B) were co-expressed in primary human T cells with the RQR8 suicide/sort marker^22^ and a first generation CD19-targeting CAR (19-ζ; Figure 1C), and were screened for their ability to resist FasL-mediated cell death and to alter T-cell activity upon binding FasL.

Upon binding FasL, Fas-CD40, Fas-DcR2, Fas-TACI, and Fas-BCMA induced NF-κB activity, where Fas-DcR2 surprisingly displayed greatest induction (Figures 1D, S1A). Fas-BCMA, Fas-CD30, and to a lesser extent Fas-CD40, exhibited constitutive NF-κB activity (Figure 1D). Fas-RANK resulted in high constitutive NF-κB activity, which paradoxically decreased in the presence of FasL (Figure 1D, S1A). Fas-CD40 induced the greatest proliferation upon binding FasL with a relative fold difference of 3.4 (Figures 1E, S1B). Fas-BCMA, Fas-HVEM, Fas-CD27, Fas-Fn14, Fas-41BB and Fas-BAFFR also induced FasL-dependent proliferation. Corresponding increases of IFNγ secretion were observed for these Fas-TNFRs (Figure S1C). Fas-LTβR and Fas-CD30 expression resulted in constitutive IFNγ secretion (Figure S1C), despite not resulting in basal or induced proliferation (Figures 1E, S1B). No IL-2 secretion was induced by any of the Fas-TNFRs in response to FasL (Figure S1D). Fas-TNFR chimeras that induced proliferation upon binding FasL (Fas-HVEM, Fas-CD27, Fas-41BB, Fas-CD40, Fas-BAFFR, Fas-BCMA and Fas-Fn14; Figure S1B) were selected for further study and assayed in response to FasL using six different donors, which confirmed the earlier findings (Figure S1E). We continued our investigations with the following chimeras: Fas-CD27, Fas-41BB, Fas-CD40, Fas-BCMA and Fas-Fn14.

### Fas-CD40 induces profound transcriptional changes

We next investigated how the Fas-TNFRs affected gene transcription using the nCounter® NanoString® platform. We first compared transcripts to 19-ζ-expressing cells which revealed two clusters of differentially expressed genes (DEGs): (1) FasΔDD, Fas-CD27 and Fas-41BB; and (2) Fas-CD40, Fas-BCMA and Fas-Fn14 (Figures 1F, 1G, S2A, e-table 1). Under basal conditions, cluster 2 induced greater gene transcription compared to cluster 1, and in the presence of FasL transcriptional increases were observed across both clusters, consistent with markers of functional activation observed above (Figures 1D, 1E, S1C). Upregulated DEGs specifically identified in cluster 2 related to: T-cell cytotoxicity; the cell cycle; chemokine and interleukin signalling; JAK-STAT, MAPK/PI3K, and NF-κB pathways; co-stimulatory molecules; T-cell memory; and oxidative phosphorylation, mitochondrial biogenesis, lipid metabolism, and glycolysis (Figures 1F, 1G, Table 1).

**Table 1.** Upregulated differentially expressed genes (DEGs) specific to Fas-CD40, Fas-Fn14 and Fas-BCMA expression. Selected list of upregulated significant (*P* < 0.05) DEGs relating to their pathway involvement. T cells were treated as described in Figure 1F.

| Gene pathway | Genes |
| --- | --- |
| T-cell cytotoxicity | *CSF2, IFNG, GZMB, PRF1, FASLG* |
| JAK-STAT signalling | *LIF, STAT1, STAT3, STAT5A, SOCS4, CRLF2* |
| MAPK/PI3K signalling | *CDC42, PIK3R1, PIK3R2, PIK3R3, RAC2, MAPK3, MAP2K2, MAP3K14* |
| Cell cycle | *NSD2, AURKA, SGO2, NEK2, MKI67, BUB1, NCAPH* |
| Oxidative phosphorylation | *NDUFA1, NDUFA2, ATP5MF, IDH3A, COX4I1, COX5B, COX6B1, COX6C, COX7A2, COX7B, COX7C* |
| Mitochondrial biogenesis | *SLC25A6, HSPE1, NAA20, COX19* |
| Lipid metabolism | *HMGCR, SCD, MID1IP1, ACACA, ACSF2* |
| Glycolysis | *PFKFB4, LDHA, PGAM1, PGK1, PKM* |
| T-cell memory | *CD45RO, SELL, LEF1, BATF3, TCF7* |
| Interleukin signalling | *EBI3, IL2RA, IL2RB, IL2RG, IL3, IL3RA, IL12RB2, IL21R, IL32, IL36A* |
| Chemokine signalling | *CCR8, CXCR3, CXCR4, CCL1, CXCL10, CXCL13, XCL1/2, CCR7* |
| NF-κB pathway | *NFKBIA, NFKB2, BCL2, BCL2L1, UBE2I, PARP1, IKBKE, RELA* |
| Co-stimulatory molecules | *TNFRSF4, TNFRSF9, TNFRSF18, ICOSLG, ICOS, CD80, CD27* |

Within cluster 2, Fas-CD40 displayed greater differential gene transcription which is exemplified by the number of uniquely transcribed genes (Figure 1H, Table 2). Notably, Fas-CD40 induced greater gene transcription for chemokine receptors: *CCR8, CXCR3*, and *CXCR4*; chemokine ligands: *CCL1* (encodes ligand for CCR8), *CXCL10* (encodes ligand for CXCR3) and *CXCL13*; and *ICOSLG* (encodes ligand for ICOS) (Figure 1H). Downregulated DEGs included inhibitory checkpoints (*CISH, LILRB3*) and pro-apoptotic markers (*BID, CASP8*) (Figures 1G, 1H, S2B). To exclude against the possibility that Fas-TNFR protein expression caused these transcriptional differences, Fas-TNFR expression varied within each cluster and was equivalent between clusters (Figure S2C).

**Table 2.**
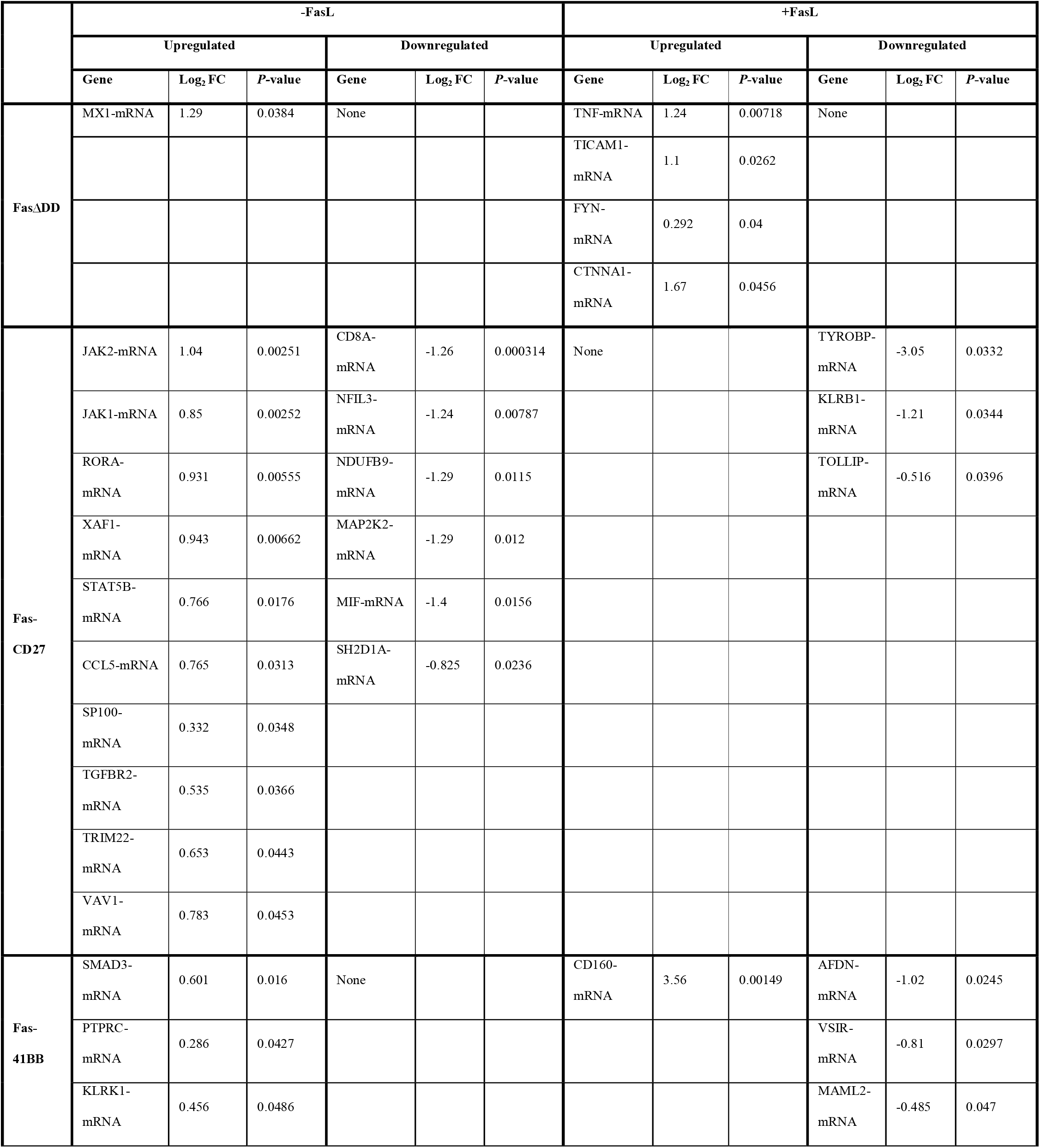
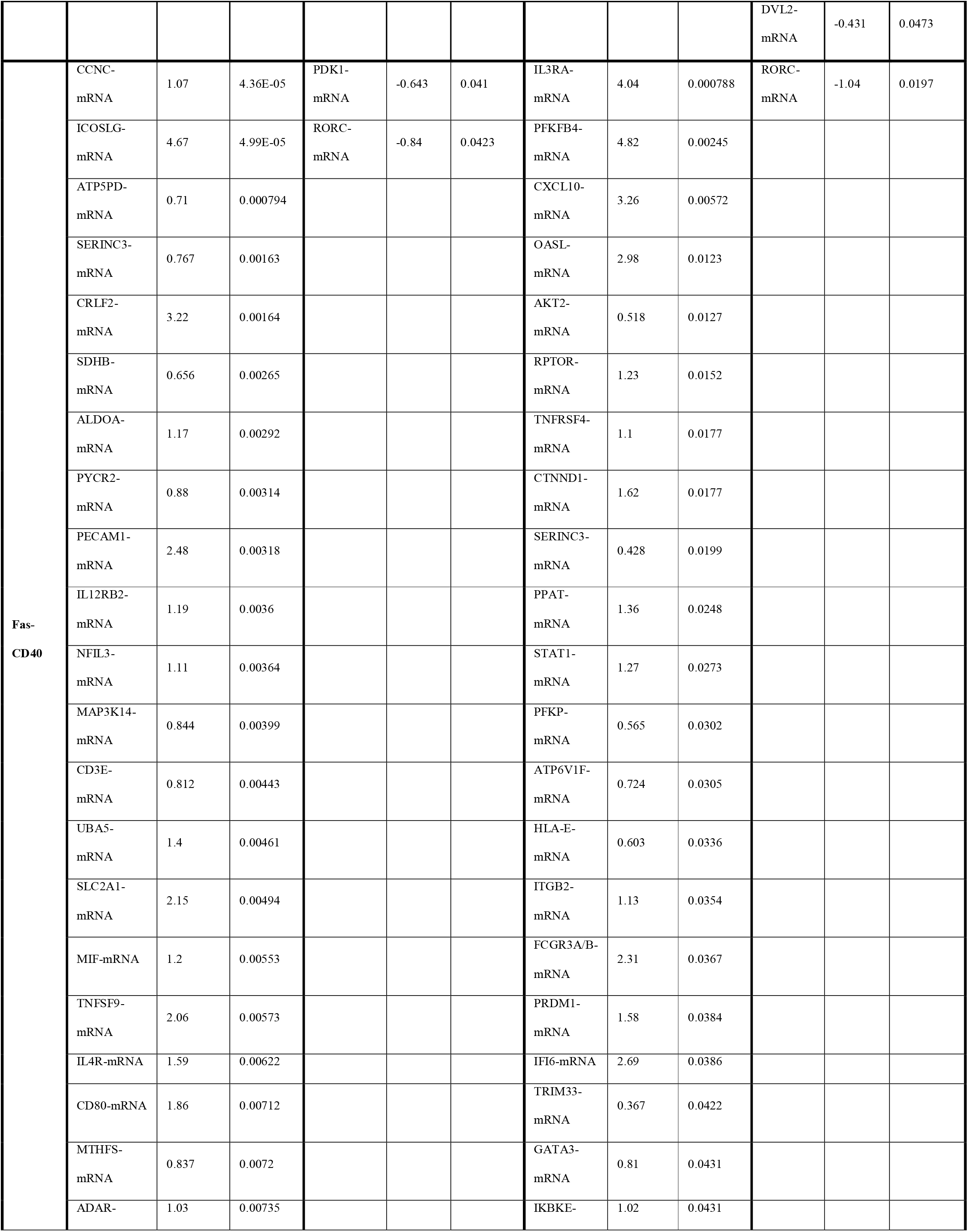
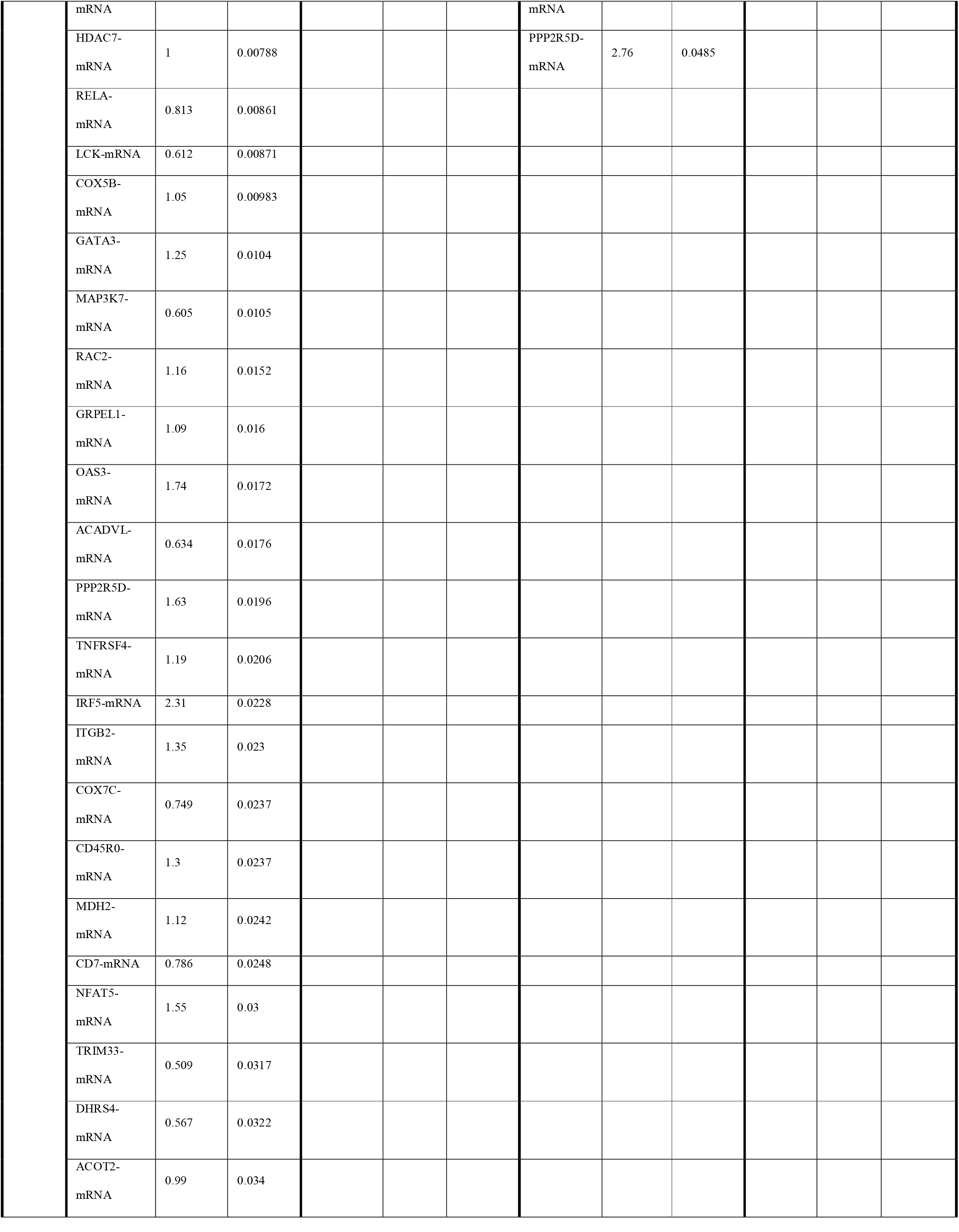
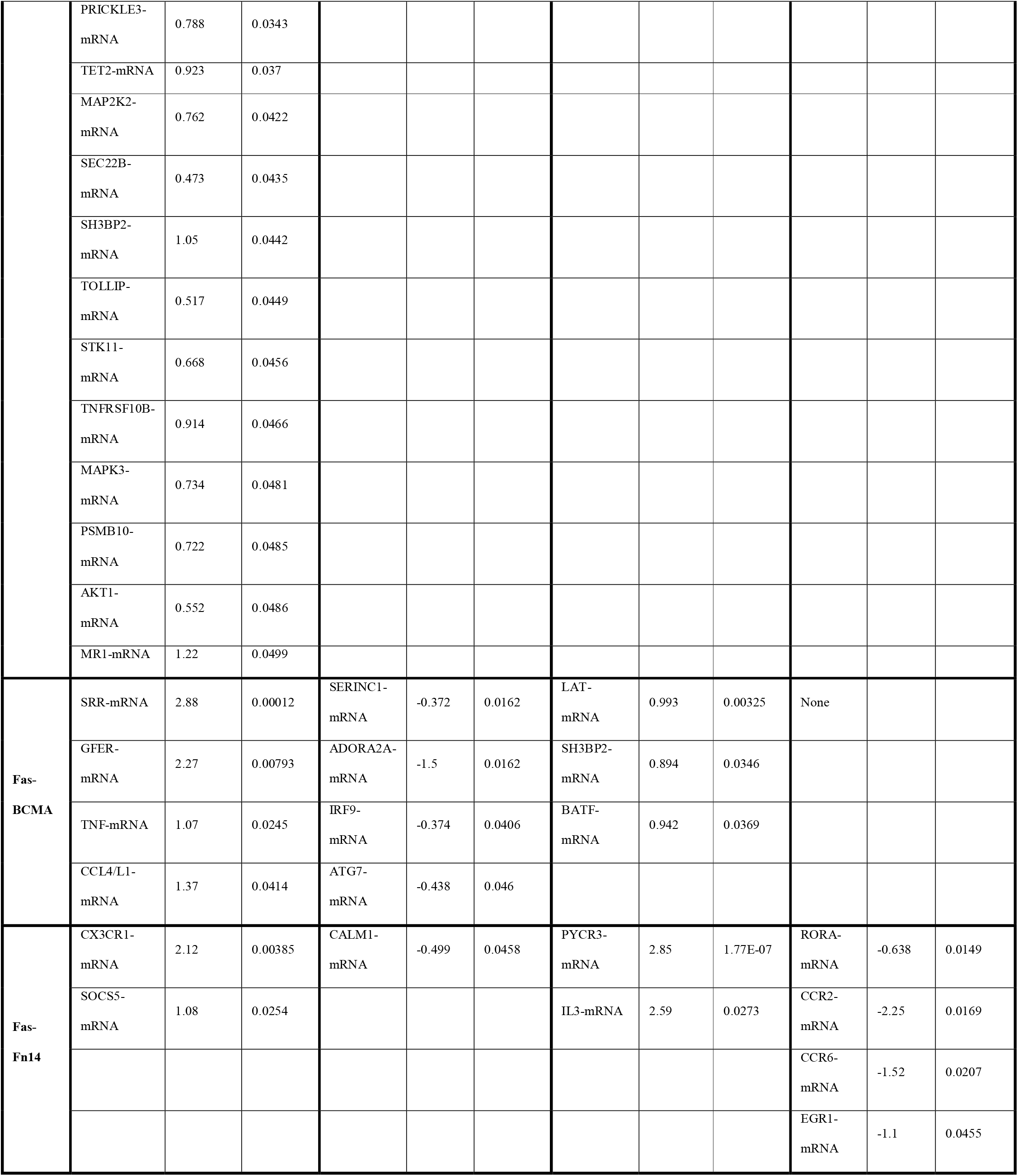
List of uniquely transcribed genes relative to 19-ζ expression. T cells were treated as described in Figure 1F. TCR diversity genes included in the nCounter® CAR-T Characterization Panel have been removed from this list, FC – fold change.

Comparative analysis of Fas-TNFRs versus FasΔDD revealed similar upregulated DEGs compared to 19-ζ such as those related to NF-κB, interleukin, and chemokine signatures (Figures 1I, S2D, e-table 2). However, upon binding FasL there were greater downregulated DEGs in cluster 2, which included markers of T-cell inhibition (*CD200, CTLA4, LILRB3, CD276, CD84*), senescence (*KLRG1*), exhaustion (*TOX*), and apoptosis (*GADD45B, CASP8, BID*).

We confirmed increased protein expression of CCR8, ICOSL and ICOS upon Fas-CD40 expression (Figure 1J), in addition to increased secretion of CCL1, CXCL10 and CXCL13 (Figure 1K), correlating with our transcriptomic analysis.

### Fas-TNFR chimeras protect from FasL-mediated kill

We next assessed how efficiently the Fas-TNFRs could rescue FasL-mediated kill. We co-expressed the Fas-TNFRs with RQR8 and a second-generation CD19-targeting CAR (19-BBζ; Figure 2A). FasΔDD, Fas-CD27 and Fas-CD40 had the highest protein expression, respectively, with Fas-41BB having the lowest (Figures 2B, 2C). Upon co-culture with SupT1 cells engineered to express FasL (Figure S3A), Fas-41BB could only partially rescue cell death as measured by cell survival, however the percentage of apoptotic cells were indistinguishable from SupT1 control cells (Figure 2D). FasΔDD and the other Fas-TNFRs fully rescued FasL-mediated cell death. Upon repeated SupT1-FasL challenge, Fas-CD40, Fas-CD27 and FasΔDD completely rescued FasL-mediated cell death, with Fas-CD40 and Fas-CD27 additionally inducing CAR T-cell proliferation (Figures 2E, 2F, S3B), consistent with the initial screen. In contrast, Fas-41BB, Fas-BCMA and Fas-Fn14 only partially rescued cell survival. Fas-TNFR protein expression correlated to 19-BBζ survival (Figure 2G). Fas-TNFR-19-BBζ CAR T cells did not proliferate autonomously when co-cultured with CD19^−^ SupT1 cells (Figure 2E), nor did they increase tonic cytotoxicity relative to 19-BBζ alone (Figure 2H). The level of cytotoxicity slightly increased against SupT1-FasL cells (Figure 2H).

**Figure 2.**
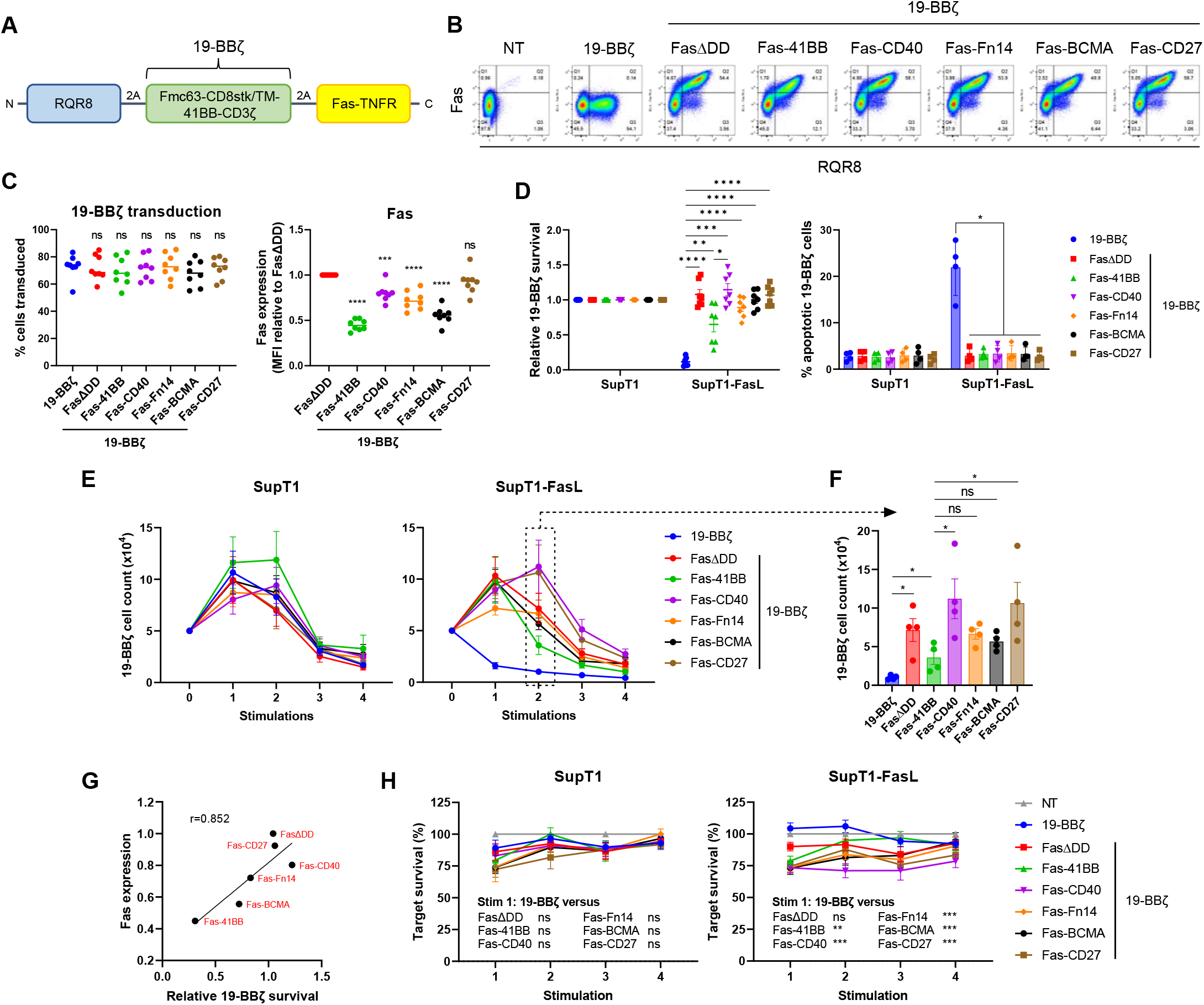
Fas-TNFRs rescue FasL-mediated kill. (A) Schematic of polycistronic transgene transduced into human T cells. 19-BBζ: Fmc63 binder fused to the endodomains of 4-1BB and CD3ζ via a CD8 stalk/transmembrane domain. (B) Representative flow cytometry plots from one human T-cell donor transduced to express either 19-BBζ alone or co-express FasΔDD or the stated Fas-TNFRs. (C) Left: transduction percentages of T cells from eight independent donors, ns – non-significant, one-way ANOVA (Dunnett’s multiple comparisons test relative to 19-BBζ). Right: median fluorescence intensity (MFI) of the Fas-TNFRs relative to FasΔDD MFI, measured from top right (Q2) quadrant in B. Eight independent donors tested, mean being shown, \*\*\**P* < 0.001, \*\*\*\**P* < 0.0001, ns – non-significant, one-way ANOVA (Dunnett’s multiple comparisons test relative to FasΔDD). (D) 19-BBζ cells were cultured with SupT1^FasKO^ or SupT1^FasKO^-FasL cells either for 72 hours (left) or five hours (right) at a 1:1 effector to target ratio (E:T), at which point 19-BBζ cell survival or percentage of apoptotic cells (Annexin V^+^ 7AAD^−^) were calculated, respectively. Seven and four independent donors were tested for the cell survival and apoptotic analysis, respectively. Error bars are SEM, \**P* < 0.05, \*\**P* < 0.01, \*\*\**P* < 0.001, \*\*\*\**P* < 0.0001, two-way ANOVA. (E) 19-BBζ cells from four independent donors were stimulated with 5×10^4^ SupT1^FasKO^ or SupT1^FasKO^-FasL cells at an initial 1:1 E:T up to four times with cell counts being analysed after each stimulation. (F) Cell numbers from E after the second round of SupT1^FasKO^-FasL stimulation. \**P* < 0.05, ns – non-significant, two-way ANOVA. (G) Mean average of relative Fas expression (from Figure 2C) *versus* mean average of relative 19-BBζ survival (from second stimulation readout in Figures 2E and S3B), r = Pearson correlation coefficient. (H) From experiment described in E, the percentage of surviving targets analysed after each target stimulation. Error bars are SEM, \*\**P* < 0.01, \*\*\**P* < 0.001, ns – non-significant, two-way ANOVA.

### Fas-CD40, Fas-BCMA, Fas-CD27 and Fas-Fn14 enhance 19-BBζ CAR efficacy

We then assessed how the Fas-TNFRs affected 19-BBζ-mediated target cell cytotoxicity by co-culturing with multiple cancer cell lines. Fas-TNFR-19-BBζ cells exhibited equivalent cytotoxicity and secretion of IFNγ and IL-2 against CD19^+^ Nalm6 and Raji cancer cells compared to 19-BBζ alone (Figures 3A, S4A). Additionally, Fas-TNFR-19-BBζ cells demonstrated equivalent cytotoxicity against SupT1 cells engineered to express the CAR cognate antigen CD19, with no background killing observed against CD19^−^ SupT1 cells (Figure S4B).

**Figure 3.**
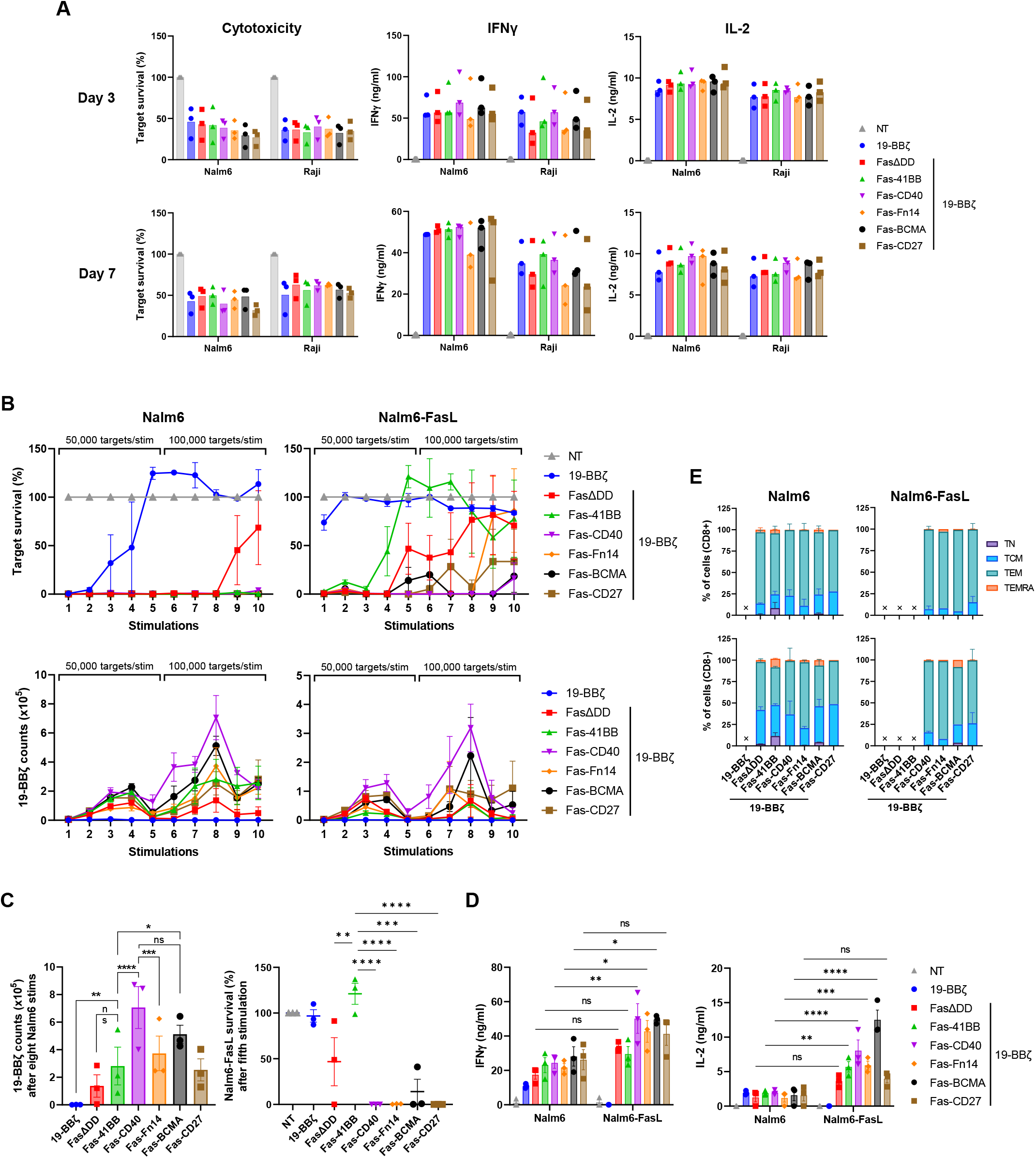
Fas-TNFRs augment 19-BBζ CAR efficacy. (A) 19-BBζ cells co-expressing FasΔDD or the Fas-TNFRs were cultured with Nalm6 or Raji cells for three or seven days at a 1:4 E:T, measuring for target survival (left), and secretion of IFNγ (middle) and IL-2 (right). Three independent donors tested, error bars are SEM. (B) 19-BBζ cells from three independent donors were stimulated up to ten times with either Nalm6^FasKO^ or Nalm6^FasKO^-FasL cells at a starting 1:8 E:T, measuring for target survival and 19-BBζ cell counts after each stimulation. Effectors were stimulated with 50,000 targets for the first five stimulations and 100,000 targets for the final five stimulations, error bars are SEM. (C) Left: 19-BBζ cell counts after eighth round of Nalm6^FasKO^ stimulation as described in B. Right: Relative target survival of Nalm6^FasKO^-FasL cells after fifth round of stimulation as described in B. \**P* < 0.05, \*\**P* < 0.01, \*\*\**P* < 0.001, \*\*\*\**P* < 0.0001, ns – non-significant, two-way ANOVA, error bars are SEM. (D) Cell culture supernatants after the first round of target stimulation from experiment described in B were analysed for IFNγ and IL-2. \**P* < 0.05, \*\**P* < 0.01, \*\*\**P* < 0.001, \*\*\*\**P* < 0.0001, ns – non-significant, two-way ANOVA, error bars are SEM. (E) T-cell memory phenotypes were analysed for CD8 (top) and CD4 (bottom) cells after the seventh stimulation from the restimulation experiment described in B. Error bars are SEM, an ‘X’ denotes where too few cells were present to accurately determine memory phenotype.

To stress-test the Fas-TNFR chimeras we set up an *in vitro* restimulation cytotoxicity assay, where 19-BBζ cells were serially challenged with Nalm6 cells and Nalm6 cells engineered to express FasL (Figure S4C). Expression of FasΔDD and the Fas-TNFRs enhanced 19-BBζ-mediated Nalm6 cytotoxicity, killing targets for all ten stimulations (Figure 3B), with Fas-CD40 inducing greatest proliferation, a 113-fold increase from the initial 1:8 effector to target (E:T) seeding ratio (Figures 3B, 3C). Fas-CD40, Fas-BCMA, Fas-CD27, and Fas-Fn14 enhanced serial cytotoxicity against Nalm6-FasL cells compared to FasΔDD, with Fas-41BB impairing serial cytotoxicity, which again correlated with CAR T-cell numbers (Figures 3B, 3C). Fas-CD40, Fas-Fn14 and Fas-BCMA induced greater IFNγ and IL-2 secretion after one round of Nalm6-FasL stimulation compared to Nalm6 cells without exogenous FasL, with Fas-41BB also secreting greater IL-2 (Figure 3D). Equivalent memory phenotypes of the 19-BBζ cells were observed for all Fas-TNFRs, adopting either a central memory or effector memory phenotype (Figure 3E).

### Fas-CD40 also enhances 19-28ζ CAR efficacy

We next investigated whether the co-stimulatory activity of the Fas-TNFRs would be affected if we replaced the co-stimulatory endodomain within the CAR from 4-1BB to CD28 (19-28ζ; Figure 4A). 19-28ζ was co-expressed in human T cells with RQR8 and the Fas-TNFRs (Figure S5A). As seen with 19-BBζ co-expression, FasΔDD displayed highest protein expression followed by Fas-CD27 and Fas-CD40, with Fas-41BB having the lowest expression (Figure 4B), which again correlated with the ability to rescue FasL-mediated cell death (Figures S5B, S5C). Fas-CD40, Fas-Fn14 and Fas-BCMA induced a slightly higher level of basal proliferation upon CD19^−^ SupT1 stimulation (Figure S5B), which correlated with increased tonic cytotoxicity, however this was not sustained (Figure S5D). This is different to that seen with 19-BBζ implying a difference in signal transduction. 19-28ζ cells co-expressing Fas-CD40, Fas-Fn14 and Fas-BCMA displayed greater cytotoxicity against SupT1-FasL cells, which was sustained with Fas-CD40 (Figure S5D).

**Figure 4.**
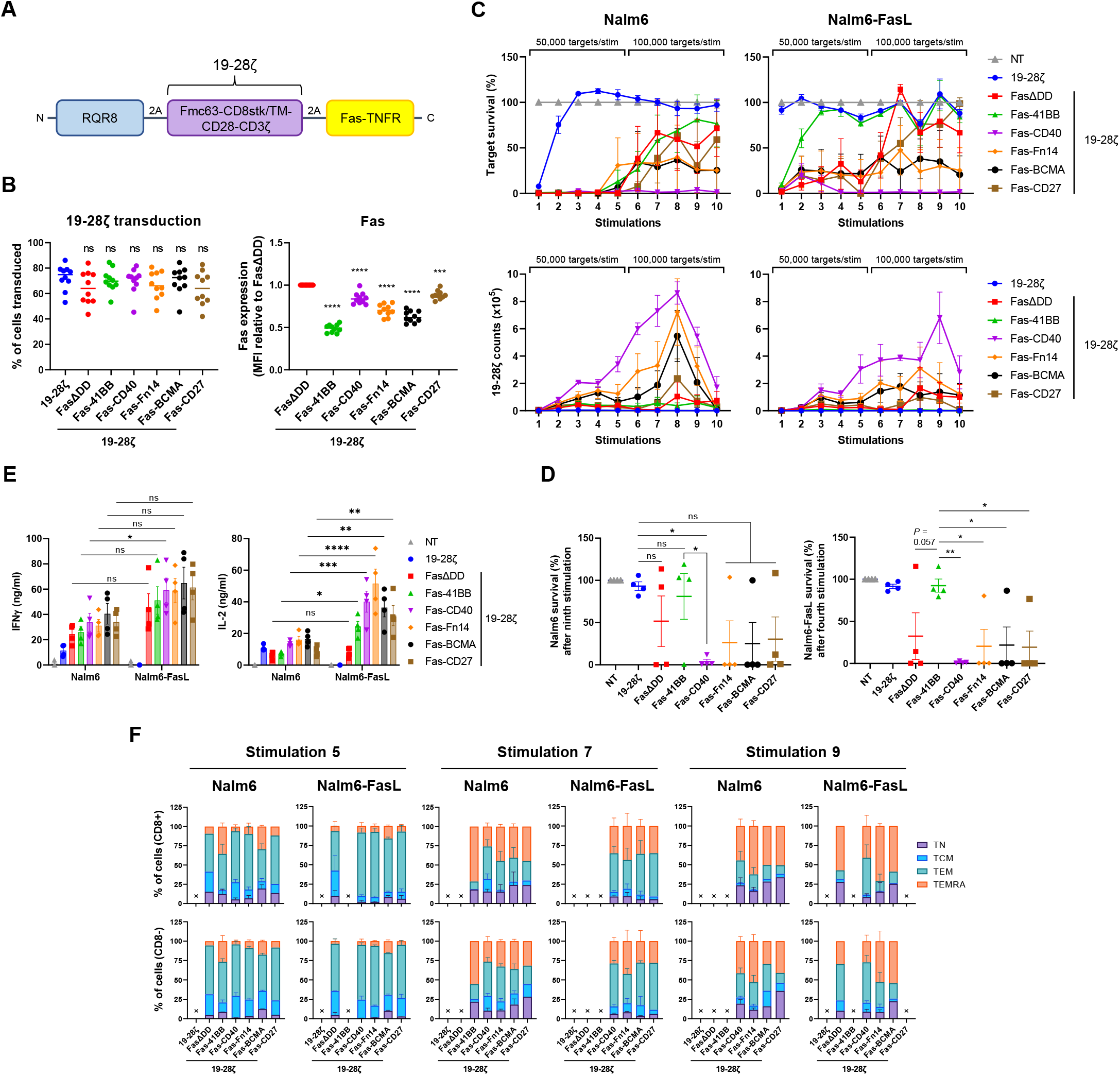
Fas-CD40 optimally enhances 19-28ζ CAR efficacy. (A) Schematic of polycistronic transgene transduced into human T cells. 19-28ζ: Fmc63 binder fused to the endodomains of CD28 and CD3ζ via a CD8 stalk/transmembrane domain. (B) Left: transduction percentages of T cells from ten independent donors, ns – non-significant, one-way ANOVA (Dunnett’s multiple comparisons test relative to 19-28ζ). Right: MFI of the Fas-TNFRs relative to FasΔDD MFI, measured from top right (Q2) quadrant in Figure S5A. Ten independent donors tested, mean being shown, \*\*\**P* < 0.001, \*\*\*\**P* < 0.0001, one-way ANOVA (Dunnett’s multiple comparisons test relative to FasΔDD). (C) 19-28ζ cells from four independent donors were stimulated up to ten times with either Nalm6^FasKO^ or Nalm6^FasKO^-FasL cells at a starting 1:8 E:T, measuring for target survival and 19-28ζ cell counts after each stimulation. Effectors were stimulated with 50,000 targets for the first five stimulations and 100,000 targets for the final five stimulations, error bars are SEM. (D) Relative target survival of Nalm6^FasKO^ (left) and Nalm6^FasKO^-FasL (right) cells after ninth or fourth round of stimulation, respectively, as described in C. \**P* < 0.05, \*\**P* < 0.01, ns – non-significant, two-way ANOVA, error bars are SEM. (E) Cell culture supernatants after the first round of target stimulation from experiment described in C were analysed for IFNγ and IL-2. \**P* < 0.05, \*\**P* < 0.01, \*\*\**P* < 0.001, \*\*\*\**P* < 0.0001, ns – non-significant, two-way ANOVA, error bars are SEM. (F) T-cell memory phenotypes were analysed for CD8 (top) and CD4 (bottom) cells after the fifth, seventh and ninth stimulations from the restimulation experiment described in C. Error bars are SEM, an ‘X’ denotes where too few cells were present to accurately determine memory phenotype.

Fas-CD40-19-28ζ cells exhibited greatest Nalm6 and Nalm6-FasL serial cytotoxicity, completely killing targets for all ten stimulations (Figures 4C, 4D), which correlated to the level of CAR T-cell proliferation (a 138-fold increase from the initial 1:8 E:T seeding ratio), and IFNγ and IL-2 secretion (Figure 4E). The proliferative capacity of Fas-CD40-19-28ζ cells was not limitless however, as CAR T-cell proliferation decreased after the last two stimulations, as did with the other Fas-TNFRs. Throughout the stimulations, Fas-CD40-19-28ζ cells maintained an earlier memory profile compared to the other Fas-TNFRs by having fewer CD45RA^+^CD62L^−^ TEMRA cells across both CD8 and CD4 populations (Figure 4F).

### Fas-CD40 optimally enhances GD2-28ζ CAR efficacy

We next investigated functionality of the Fas-TNFRs in the context of a CAR targeting a different antigen, the disialoganglioside GD2, to confirm applicability across multiple CAR architectures. Fas-TNFRs (located at the N-terminus) were co-expressed with RQR8 and a GD2-targeting CAR (GD2-28ζ; Figures S6A, S6B). Fas-CD27, Fas-CD40 and FasΔDD had highest protein expression (Figure S6C), with all Fas-TNFRs rescuing FasL-mediated apoptosis (Figure S6D). Without target stimulation all Fas-TNFR-GD2-28ζ cells had equivalent memory phenotypes and exhaustion-associated marker expression (Figure S6E) and exhibited equivalent cytotoxicity against SupT1 cells engineered to express GD2 (Figure S6F). Upon serial target stimulation, Fas-CD40 optimally enhanced GD2-28ζ- mediated cytotoxicity against SupT1-GD2 and SupT1-GD2-FasL cells, which correlated with the level of CAR T-cell proliferation (Figures S6G, S6H), similar to that observed with 19-28ζ, however with the effects less pronounced. Upon serial target stimulation, Fas-CD40-GD2-28ζ cells trended to have an earlier memory phenotype compared to GD2-28ζ or the other Fas-TNFRs, as seen by fewer TEMRA and greater effector memory cells (Figure S6I), and CD4^+^ Fas-CD40-GD2-28ζ cells expressed fewer exhaustion-associated markers (Figure S6J).

### CAR T-cell derived FasL a major source for Fas-TNFR activation

We observed from the *in vitro* restimulation experiments that co-expression of the Fas-TNFRs augmented both CD19-CAR and GD2-CAR T-cell proliferation against target cells not exogenously expressing FasL (Figures 3B, 4C, S6G). Staining for surface FasL expression in SupT1 and Nalm6 cells revealed that they did not express FasL, even in the presence of IFNγ (Figures 5A, S7). Moreover, SupT1 cells cultured with 19-BBζ or GD2-28ζ cells did not induce FasL-mediated CAR T-cell apoptosis, further evidence that SupT1 cells did not express FasL (Figure 5B). This indicated that tumour derived FasL was not the major source of FasL in this co-culture. We therefore hypothesised that T-cell derived FasL upregulated upon activation could be responsible. To address this, Fas-CD40-GD2-28ζ cells were serially stimulated with an anti-CAR idiotype antibody in the presence of anti-Fas and anti-FasL blocking antibodies. As expected, addition of anti-Fas/FasL antibodies significantly decreased Fas-CD40-GD2-28ζ cell proliferation upon CAR activation, with CAR T-cell FasL upregulation confirmed (Figure 5C).

**Figure 5.**
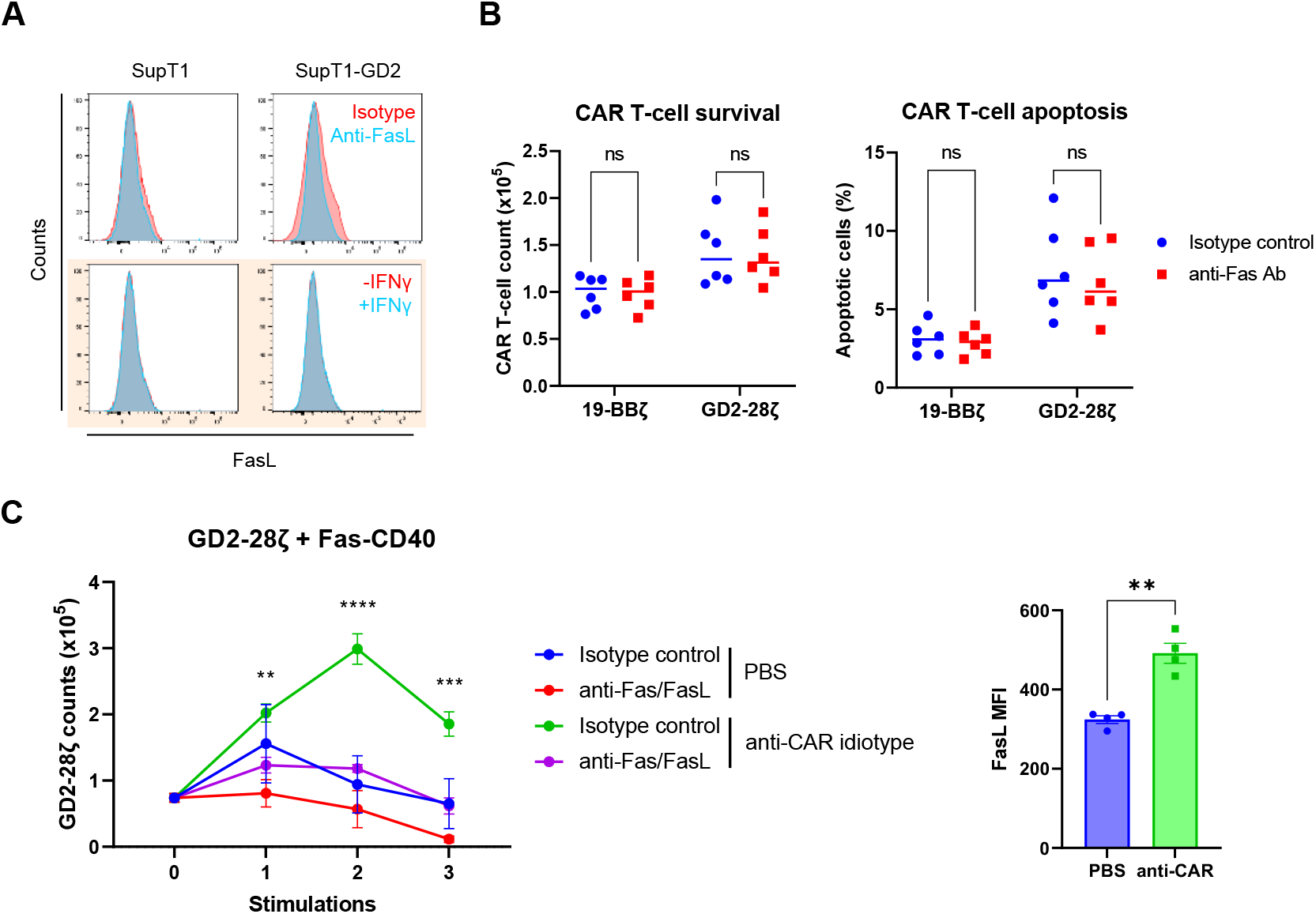
Fas-CD40 enhances CAR T-cell proliferation without exogenous source of FasL. (A) Top: SupT1 and SupT1-GD2 (Fas^+/+^) cells were surface stained with an anti-FasL antibody or an isotype control antibody. Bottom (in coloured box): SupT1 or SupT1-GD2 cells were treated with vehicle (PBS) or IFNγ (100 ng/ml) for 20 hours, and then analysed for FasL surface expression. (B) 1×10^5^ 19-BBζ or GD2-28ζ cells were cultured with SupT1 cells at a 1:1 E:T in the presence of either an isotype control or anti-Fas blocking antibody (1 μg/ml) for either 72 hours (left) or 5 hours (right), at which point CAR T-cell survival or percentage of apoptotic cells (Annexin V^+^ 7AAD^−^) were analysed, respectively. Six independent donors tested, mean being shown, ns – non-significant, two-way ANOVA. (C) Left: GD2-28ζ cells co-expressing Fas-CD40 were either unstimulated (PBS) or stimulated three times with an immobilised anti-CAR idiotype (anti-Huk666) antibody (1 μg/ml) in the presence of either an isotype control or anti-Fas and anti-FasL antibodies (1 μg/ml per antibody), where CAR T-cell counts were measured after each stimulation (counts measured four days after each stimulation). Three independent donors tested, error bars are SEM, \*\**P* < 0.01, \*\*\**P* < 0.001, \*\*\*\**P* < 0.0001, two-way ANOVA (statistics comparing isotype control *versus* anti-Fas/FasL condition upon CAR stimulation). Right: FasL expression of Fas-CD40-GD2-28ζ cells after first round of anti-CAR stimulation. \*\**P* < 0.01, two-tailed paired *t*-test, error bars are SEM.

### Fas-CD40 significantly enhances 19-BBζ-mediated anti-tumour responses in vivo

From our *in vitro* restimulation experiments with 19-BBζ it was not clear which Fas-TNFR provided the greatest co-stimulatory advantage, as Fas-CD40, Fas-BCMA and Fas-CD27 all exhibited similar efficacies (Figure 3B). Therefore, we continued our investigations *in vivo* using the xenograft Nalm6 model in NOD-*scid-*IL2Rgamma^null^ (NSG) mice. T cells from two human donors were transduced to express 19-BBζ alone or co-express FasΔDD or the Fas-TNFRs (Figure 6A). Co-expression of Fas-CD40 displayed greatest tumour killing out of all the Fas-TNFRs, with Fas-CD40-19-BBζ treatment having significantly lower tumour burden compared to Fas-41BB-19-BBζ (*P* = 0.0295) (Figures 6B, 6C). Furthermore, co-expression of Fas-CD40 and Fas-CD27 significantly improved mouse survival relative to Fas-41BB (*P* = 0.0145 and *P* = 0.0228, respectively) (Figure 6D). There was no significant survival advantage between FasΔDD and Fas-41BB treatments.

**Figure 6.**
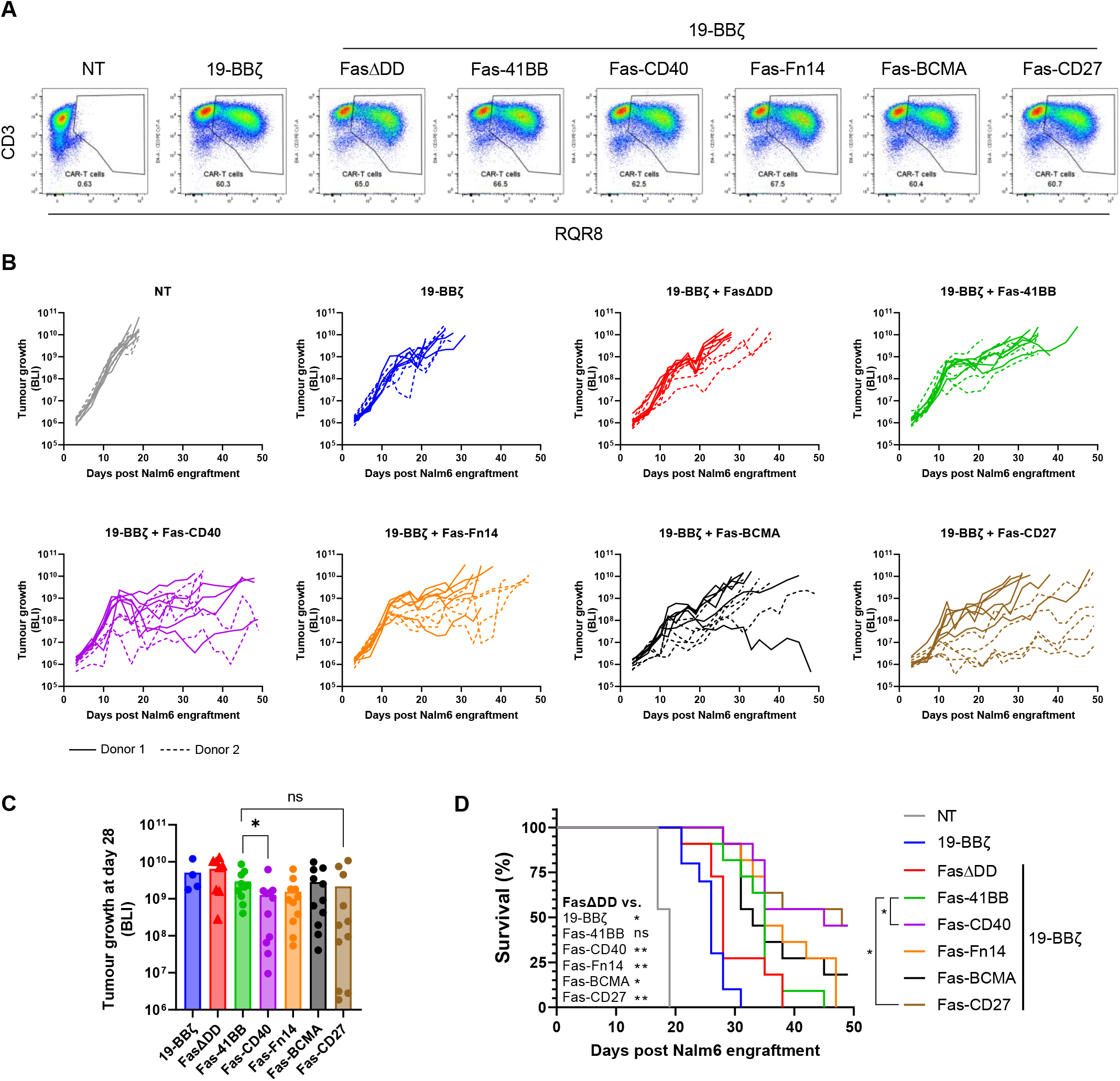
Fas-CD40 is superior for enhancing 19-BBζ-mediated anti-tumour killing and survival *in vivo*. (A) Representative flow cytometry plots from one human donor displaying transduction percentages of 19-BBζ cells at day of intravenous (iv) injection. (B) NSG mice were administered with 0.5×10^6^ Nalm6 cells expressing firefly luciferase (FLuc) by iv injection, engrafted for four days, and then 3×10^6^ CAR T cells (or equivalent total NT cells) were administered by iv injection in the tail vein (*n* = 11 mice per cohort, pooled from two independent studies). Tumour growth was measured three times weekly by bioluminescence (BLI) readout. Individual mouse tumour growths per cohort being shown. (C) Tumour growths from B at day 28 post Nalm6 engraftment. Mann-Whitney *U* test (two-tailed), \**P* < 0.05, ns – non-significant. (D) Kaplan-Meier curve from data shown in B showing overall mouse survival. Mantel-Cox test, \**P* < 0.05, \*\**P* < 0.01, ns – non-significant.

## DISCUSSION

The Fas receptor is ubiquitously expressed in T cells and its activation upon binding FasL triggers apoptosis^4–7^. Many cancer cells express FasL, in addition to TME cells such as MDSCs, CAFs, Tregs and the tumour endothelium^5,6^, as well as T cells themselves^10,11^. Therefore, the Fas/FasL checkpoint may inhibit cancer immunotherapeutic approaches such as adoptive cell therapy by limiting the persistence of T cells.

Strategies to overcome the Fas/FasL checkpoint include systemic antibody blockade^12–14^. Approaches applicable to adoptive immunotherapy also include genetic manipulation by *FAS* knockdown and knockout using siRNA and CRISPR/Cas9, respectively^15,16^. Additional described approaches include the expression of non-functional Fas, such as FasΔDD and Fas-41BB. Both FasΔDD and Fas-41BB rescue T cells from FasL-mediated kill, where the Fas-41BB chimera has the additional advantage of transmitting co-stimulatory signals upon FasL binding^5,17,18^.

Different TNFRs may transmit qualitatively different signals due to alternative TRAF recruitment to the TNFR. We hypothesised that chimeric Fas-TNFRs that had a different TNFR endodomain to 4-1BB might have different biological effects and hence may be able to better augment immunotherapeutic approaches like CAR T cells. We generated a library of 17 Fas-TNFR chimeras, identifying Fas-CD40, Fas-CD27, Fas-BCMA, Fas-Fn14, Fas-41BB, Fas-HVEM and Fas-BAFFR chimeras that could induce T-cell proliferation upon binding FasL, with Fas-CD40 eliciting greatest proliferation.

Transcriptional profiling of the five most functional chimeras identified two clusters: (1) Fas-CD27 and Fas-41BB; and (2) Fas-CD40, Fas-BCMA and Fas-Fn14, with cluster 2 displaying greater upregulated DEGs relating to the cell cycle; chemokine and interleukin signalling; JAK-STAT, MAPK/PI3K, and NF-κB pathways; and metabolism. Notably, Fas-CD40 upregulated chemokine receptor/ligand genes: *CCR8, CXCR3, CXCR4, CCL1, CXCL10*, and *CXCL13*; which were confirmed at the protein level and have all been implicated in T-cell trafficking and could facilitate T-cell homing to tumours^23–26^. Interestingly, CCR8 overexpression in CAR T cells enhanced tumour-homing, driven by a feed-forward loop of activated CAR T cells secreting CCL1 (the cognate ligand for CCR8)^23^. Expression of the Fas-TNFRs increased 19-BBζ-mediated *in vitro* serial cytotoxicity over FasΔDD, except for Fas-41BB against FasL-expressing targets, with Fas-CD40 inducing greatest proliferation upon serial target stimulation. Further, we showed Fas-CD40, Fas-Fn14, Fas-BCMA and Fas-CD27 enhanced 19-BBζ efficacy *in vivo* compared to FasΔDD, with Fas-CD40 demonstrating a significant benefit over Fas-41BB. There was no significant survival advantage *in vivo* between FasΔDD and Fas-41BB, suggesting that complementary trans-acting signalling domains between chimeras enhances CAR T-cell efficacy, rather than increasing the amplitude of one signalling pathway. Incorporation of trans-acting chimeric receptors/signalling domains to enhance CAR T-cell activity has been previously reported^27–29^.

Enhanced CAR T-cell mediated serial cytotoxicity and proliferation upon Fas-CD40, Fas-BCMA and Fas-Fn14 expression were confirmed in the context of a CD28-containing CAR (19-28ζ) and a CAR targeting a different cognate antigen (GD2-28ζ). Fas-CD40 coupled with 19-28ζ exhibited greater tonic activity compared to 19-BBζ, suggesting crosstalk between signalling pathways. One explanation could be that increased CAR tonic signalling mediated by CD28^30,31^, induces greater FasL upregulation, triggering a feed-forward loop via Fas-CD40:FasL paracrine interactions. Indeed, Künkele and colleagues demonstrated that CD28-containing CARs caused activation-induced cell death via upregulated CAR T cell-derived FasL^10^. Importantly though, this tonic activity did not persist, and rather than this tonic activity induce functional exhaustion/dysfunction, the opposite was true with Fas-CD40 maintaining the capacity for serial target killing. Importantly, we did not observe any evidence of autonomous proliferation with Fas-CD40, or with any other Fas-TNFR, co-expressed with either 4-1BB- or CD28-containing CARs.

Interestingly, we observed the Fas-TNFRs enhanced CAR T-cell proliferation and anti-tumour cytotoxicity even when we did not enforce FasL expression on target cells. We subsequently demonstrated that a major source of FasL for Fas-TNFR activation derives from CAR T cells themselves, an effect observed with TCR-engineered-Fas-41BB cells^17,18^. Expression of the Fas-TNFRs therefore creates a self-regulatable way to augment CAR T-cell activation, irrespective of tumour FasL expression, whereby CAR activation (signals one and two) upregulates FasL surface expression, binding the Fas-TNFR on a sister CAR T-cell, which delivers an additional third signal to the CAR T-cell (Figure 7). This is akin to physiological TCR-mediated activation between a T-cell and an APC, with APCs delivering additional signals to T cells via presentation of TNFR ligands, a concept explored with expression of full length 4-1BB or OX40 in CAR T-cells which enhanced their efficacy^32,33^.

**Figure 7.**
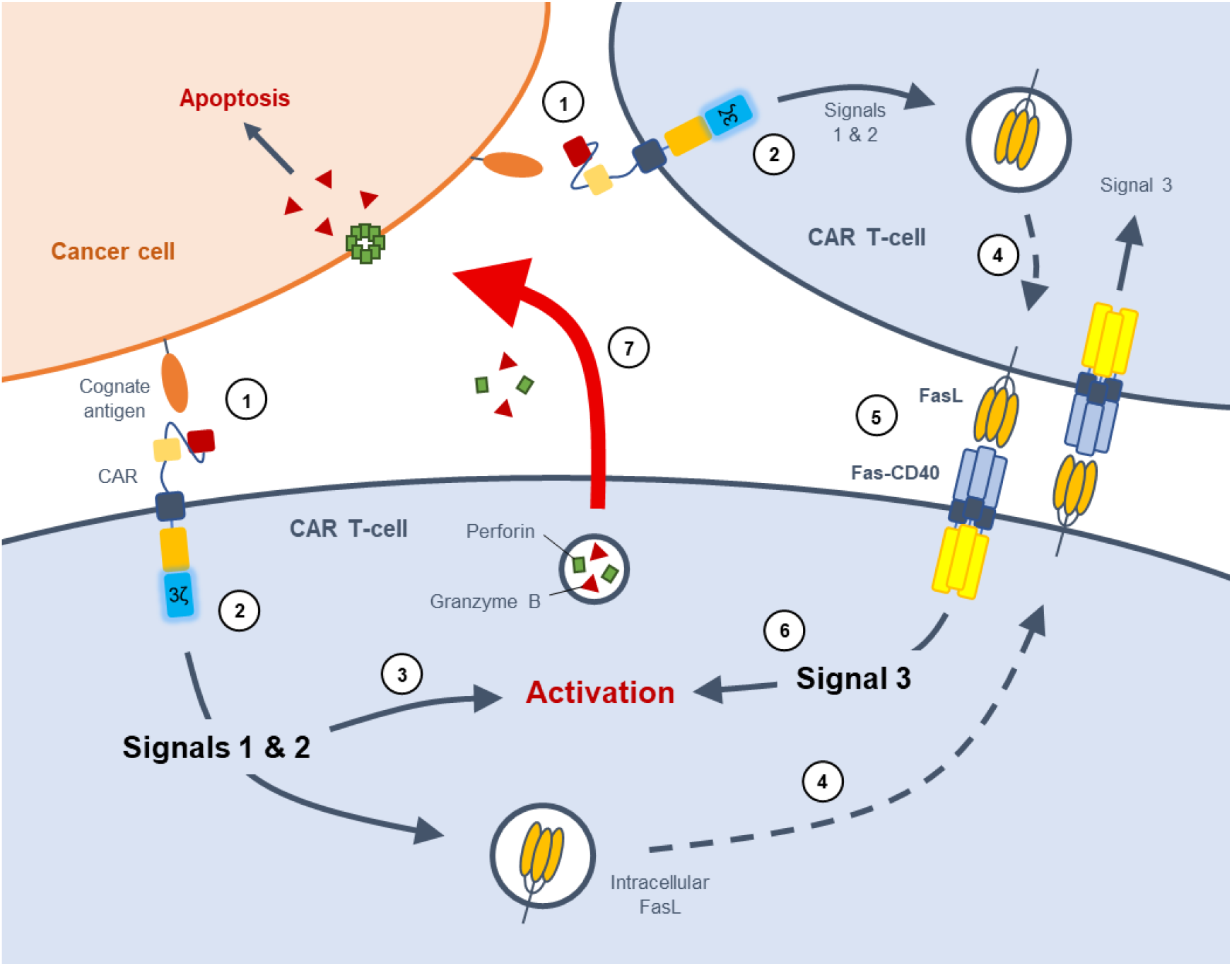
Universal application of Fas-TNFRs to enhance CAR T-cell therapy. Cartoon illustrating how Fas-TNFRs augment CAR T-cell efficacy. The CAR binds the cognate antigen on the cancer cell forming an immunological synapse (1). The co-stimulatory domain and CD3ζ within the CAR undergo signal transduction delivering signals 1 and 2 to the cell (2), leading to CAR T-cell activation (3). Intracellular stores of FasL are then trafficked to the plasma membrane (4), where Fas-TNFR binds to upregulated FasL on neighbouring CAR T cells (5), delivering signal 3 to the cell, augmenting CAR T-cell activation (6). Upon activation, the CAR T-cell induces target cell killing (7).

As well as the chimeras highlighted above, some chimeras exhibited different effects on T-cell function. Fas-LTβR and Fas-CD30 induced constitutive IFNγ secretion, however they could not rescue FasL-mediated kill. Expression of full-length LTβR in T cells has been shown to potentiate TCR-activated IFNγ secretion^34,35^, however it did not constitutively induce IFNγ, therefore the constitutive IFNγ secretion observed with Fas-LTβR suggests the Fas ecto- and transmembrane domains might be clustering the chimera to form dimers, as has been previously described for Fas prior to ligand-binding^36^. Fas-RANK induced strong constitutive activation of NF-κB, an effect observed with overexpressing full-length RANK^37^, however this did not correlate to an enhancement of proliferation or IFNγ secretion. Fas-DcR2 remarkably induced very high levels of NF-κB activation upon binding FasL, consistent with the literature that DcR2 activates NF-κB^21^, however this did not correlate to increased proliferation or IFNγ release.

Differences in functional activity between Fas-TNFRs are likely due to qualitative differences in TRAF recruitment. For example, CD40 and BCMA recruits TRAFs 1-3, 5 and 6 upon activation; whereas 4-1BB only recruits TRAFs 1-3^8^. However, this cannot solely explain the differences in Fas-TNFR performance, as OX40 and RANK, which did not induce proliferation upon binding FasL, also interacts with TRAFs 1-3, 5 and 6^8^. Quantitative differences in the amount of recruited TRAFs to each TNFR will likely also dictate the amplitude of signalling output. TRAF6 could be the likely determining molecular factor differentiating between Fas-TNFR function, as TRAF6 appears to be the predominant TRAF for CD40-induced NF-κB activation in dendritic cells^38^, and that TRAF6 also binds BCMA and Fn14, the chimeras of which augmented CAR T-cell activity akin to Fas-CD40. TRAF6 is unique from the other TRAFs by: having a different binding motif (P-x-E-x-x-[acidic/aromatic residue]); being involved beyond TNFR signalling such as IL-1R and toll-like receptor signalling^39^; and being able to activate the Src-family tyrosine kinases resulting in Akt activation via phosphoinositide 3-kinase (PI3K), in addition to activation of transcription factors NF- κB and AP-1, the latter of which being common amongst other TRAFs^39,40^. Fas-TNFR co-expression with 4-1BB- or CD28-containing CARs adds a further layer of signalling complexity, particularly because CD28 belongs to the immunoglobulin superfamily and as such recruits different signalling proteins to TNFRs, namely SH2- and SH3-domain containing proteins such as Grb2, PI3K, and Lck^41–44^.

Protein expression of the Fas-TNFR chimera does not appear to determine its co-stimulatory activity, as Fas-CD27 consistently had the highest expression across multiple CAR architectures, however its ability to augment CAR T-cell activation varied depending on the CARs co-stimulatory domain. Similarly, Fas-BCMA, which had relatively low expression, was able to enhance CAR T-cell activation akin to Fas-CD40 with either 4-1BB- or CD28-containing CARs. It is possible that even a low level of Fas-TNFR expression will saturate the amount of available endogenous TRAFs. Fas-TNFR expression does seem particularly important for rescuing FasL-mediated kill, however.

CD40 is typically expressed in antigen presenting cells (APCs) such as macrophages, B cells and dendritic cells, interacting with CD40 ligand on T cells, functioning in a co-stimulatory manner known to activate both canonical and non-canonical NF-κB pathways^45,46^. However, CD40 is also expressed in T cells, similarly functioning in a co-stimulatory manner: activating canonical and non-canonical NF-κB pathways, AP-1 and the AP-1 activator JNK^47^; generating T-cell memory and ameliorating exhaustion^48,49^. CD40 has been identified in several independent screens for enhancing T-cell function^35,50^; and has been synthetically incorporated into CAR T cells, either as a separate module or incorporated into the CAR architecture, displaying superior anti-tumour activity compared to conventional CAR T cells, facilitated by enhanced proliferation and maintaining T-cell stemness/memory^29,51–56^. BCMA is expressed in mature B lymphocytes and has been synthetically expressed in CAR T cells, augmenting proliferation^56^; whereas Fn14, expressed in healthy tissue and particularly in solid tumours such as glioblastoma^57^, has not been previously synthetically expressed in T cells to alter their function. CD27 is a well-known T-cell co-stimulatory protein, enhancing T-cell function and generating memory^35,58–60^. CD27 has been incorporated into CARs, displaying equivalent *in vivo* functionality to 4-1BB and CD28^61,62^.

We have extended possibilities of engineering T cells to be resistant to FasL-mediated apoptosis by showing that chimeras of Fas with a range of TNFR endodomains can have potentially useful biological functions. Fusion proteins such as Fas-CD40 may enhance anti-tumour activity when co-expressed with CARs.

## MATERIALS AND METHODS

### Cell lines

HEK-293T (ATCC; CRL-11268) cells were cultured in Iscove’s Modified Dulbecco’s Medium (Sigma; I3390) supplemented with 10% FBS (Biosera; FB-1058) and 2 mM GlutaMAX™-1 (Gibco; 35050). SupT1 (ECACC; 95013123), Nalm6 (DSMZ; ACC 128), Raji (ECACC; 85011429) and Jurkat E6.1 (ECACC; 88042803) cell lines were cultured in RPMI-1640 Medium (Sigma; R0883) supplemented with 10% FBS and 2 mM GlutaMAX. Cell lines were cultured at 37°C, 5% CO_2_.

### DNA construct generation

All open reading frames were cloned into the MoMLV-based retroviral genome construct SFG. Linear DNA fragments (gBlocks), encoding codon optimised open reading frames (GeneArt), were synthesised (IDT) and amplified using Q5 DNA polymerase (NEB; M0491L) and oligonucleotide primers (IDT). The resulting PCR products were fractionated on an agarose gel, purified using the QIAquick gel extraction kit (Qiagen; 28706), and digested with *Esp3 I* or *Bsa I*-HF v2 (NEB; R0734L and R3733L, respectively). Digested DNA fragments were purified using the QIAquick PCR purification kit (Qiagen; 28106) and ligated to gel-purified plasmid backbones using T4 DNA ligase (Roche; 10799009001). New England Biolabs high efficiency competent 5α *E. coli* (NEB; C2987U) were transformed with the ligation reactions, plated onto LB agar containing ampicillin (final concentration of 100 μg/mL) and incubated overnight at 37°C.

Where more than one open reading frame was inserted into the retroviral genome plasmid, self-cleaving peptide sequences derived from *Thosea asigna* virus 2A (T2A), equine rhinitis A virus polyprotein (E2A) or porcine teschovirus-1 2A (P2A) were introduced to facilitate expression from a single mRNA transcript. The suicide/sort marker RQR8 was included in the retroviral genome plasmids to enable detection of transduced cells^22^.

### Retroviral production

1.5×10^6^ HEK-293T cells were transiently transfected with an RD114 envelope expression plasmid (RDF, a gift from M. Collins, University College London), a Gag-pol expression plasmid (PeqPam-env, a gift from E. Vanin, Baylor College of Medicine), and the transgene of interest expressed in a retroviral (SFG) vector plasmid at a ratio of 1:1.5:1.5 (total DNA=12.5 μg). Transfections were performed with GeneJuice® (Millipore; 70967) according to the manufacturer’s instructions and viral supernatants were harvested 48 hours post transfection and stored at -80°C.

### Calculation of functional retroviral titres

Functional viral titres of retroviral supernatant were calculated using frozen supernatant on primary human T cells activated with TransAct (Miltenyi Biotec; 130-111-160), 10 ng/ml IL-7 (Miltenyi Biotec; 130-095-367) and 10 ng/ml IL-15 (Miltenyi Biotec; 130-095-760) for 48 hours. Retroviral supernatant was serially diluted into 24-well tissue culture plates (Corning; 351147) coated with RetroNectin (Takara Bio; T100B), where 3×10^5^ activated T cells were seeded and then spun by centrifugation for 1000 *g*, 40 minutes at room temperature, and then cultured at 37°C, 5% CO_2_. Transduced cells were identified by measuring for CAR expression using anti-fmc63 and anti-Huk666 idiotypes (both produced in house). Viral titres were calculated with T cells that were less than 20% transduced.

### Transduction of primary human T cells and cancer cell lines

Peripheral blood mononuclear cells (PBMCs) were isolated from whole human blood (NHS Blood and Transplant) by density centrifugation with Ficoll-Paque Plus (GE-Healthcare; GE17-1440-03) according to the manufacturer’s instructions. Isolated PBMCs were activated with TransAct and 10 ng/ml IL-7 and IL-15 and cultured in RPMI-1640 supplemented with 10% FBS and 2 mM GlutaMAX and cultured at 37°C, 5% CO_2_. 48 hours post activation, 1×10^6^ PBMCs were seeded onto RetroNectin-coated 6-well tissue culture plates (Corning; 351146) with retroviral vector and spun by centrifugation for 1000 *g*, 40 minutes at room temperature, and then cultured at 37°C, 5% CO_2_. PBMCs were transduced at equal multiplicity of infections (MOIs) across all cohorts. SupT1 and Nalm6 cell lines were transduced in a similar manner to PBMCs, without activation with TransAct and IL-7/IL-15 and using non-titrated retroviral supernatant. Expression of the transgene in PBMCs and cancer cell lines were assessed 72 hours post transduction by flow cytometry. RQR8 expression was detected using an anti-CD34 antibody.

### Flow cytometry and antibodies

Flow cytometry was performed using MACSQuant 10 and X flow cytometers (Miltenyi Biotec). All staining, unless specified otherwise, was performed at room temperature for 10 minutes, protected from light, with antibodies diluted in either PBS (Sigma; D8537) or cell staining buffer (Biolegend; 420201), or otherwise specified. Cell viability dyes used were 7-AAD (Biolegend; 420404) or Sytox blue (Fisher Scientific; S34857). Antibodies were from Biolegend, unless otherwise stated. Antibodies used were: CD2-PE (300208), CD3-PE Cy7 (344816), CD34-APC (R&D Systems; FAB7227A), GD2-APC (357306), FasL-BV421 (306412), Fas-PE (305608), Fas-APC Cy7 (305636), CCR8-PE (360604), ICOSL-PE (309404), ICOS-PE (313508), CD45RA-PE Texas Red (Invitrogen; MHCD45RA17), CD62L-Pacific blue (304826), LAG3-FITC (369308), PD-1-PE (329906), TIM3-BV421 (345008), and CD8-APC Cy7 (301016). Anti-CD19 and anti-GD2 CARs were detected using anti-Fmc63 and anti-Huk666 idiotypes, respectively (produced in-house), and anti-mouse IgG secondary antibody conjugated to Alexa Fluor 647 (Jackson ImmunoResearch; 115-605-071).

### Generation of antigen-expressing cell lines and reporter cell lines

For the generation of FasL-expressing cell lines, SupT1 and Nalm6 cell lines were nucleofected (Lonza) with Cas9 ribonucleic protein (RNP) complexes in SF buffer (Lonza), using the pulse codes CM-150 or CV-104, respectively. RNP complexes were formed using 50 pmol of Alt-R® S.p. HiFi Cas9 Nuclease V3endonuclease (IDT; 1081060) and 100 pmol of the following single guide RNAs ([sgRNAs]; Synthego) targeting the human *FAS* locus; sgRNA 1: ggaguugaugucagucacuu; sgRNA 2: gugacugacaucaacuccaa; sgRNA 3: ugacaucaacuccaagggau; sgRNA 4: cuuccucaauuccaaucccu. Knockout (KO) efficiency was determined by flow cytometry, staining for Fas expression. Non-electroporated Fas^+^ cells were eliminated after addition of 100 ng/ml *Mega*FasL (AdipoGen; AG-40B-0130-3010) for 48 hours. SupT1^FasKO^ and Nalm6^FasKO^ cells were then transduced with retroviral supernatant to express human FasL, where transduction efficiency was measured by flow cytometry staining for FasL.

To produce SupT1 cells to express human CD19, SupT1 cells were transduced with retroviral supernatant to express human CD19 and were sorted for CD19 expression by fluorescence-activated cell sorting (FACS) on the BD FACSMelody™ Cell Sorter according to the manufacturer’s instructions. To produce SupT1 cells to express human GD2, wild-type SupT1 or SupT1^FasKO^ cells were transduced with retroviral supernatant to express GD2- and GD3-synthases, separated by a 2A self-cleaving peptide, and were then sorted for GD2 expression by FACS on the BD FACSMelody™ Cell Sorter. To produce SupT1 cells expressing GD2 and/or FasL for the GD2-28ζ *in vitro* restimulation experiments in Figure 5, SupT1^FasKO^ cells were transduced with a retroviral supernatant to express GD2- and GD3-synthases (for GD2 alone), or retroviral supernatant to express FasL, or a dual-transduction with retroviral supernatants for both GD2/GD3 synthases and FasL. SupT1 cells were then sorted for GD2 and/or FasL expression by FACS on the BD FACSMelody™ Cell Sorter.

To produce the NF-κB Jurkat reporter cell line, Jurkat E6.1 cells were electroporated by nucleofection (using platform from Lonza) with a plasmid encoding five copies of an NF-κB response element, luciferase (Promega; N1111), and a hygromycin resistance gene. Jurkat cells expressing the NF-κB reporter were cultured under hygromycin selection (100 μg/ml).

### *In vitro* cytotoxicity and proliferation assays

CAR T cells were co-cultured with 5×10^4^ target cells (unless stated otherwise) at the stated E:T, where target cells were detected by flow cytometry by the absence of CD2, CD3 and RQR8 expression. Surviving target cells were made relative to surviving targets cells treated with non-transduced (NT) T cells. The number of CAR T cells were quantified using CountBright™ Counting Beads (ThermoFisher; C36995). For the restimulation experiments, CAR T cells were initially co-cultured with 5×10^4^ target cells at the stated E:T, and then restimulated with target cells as described, twice weekly for up to a total of ten target stimulations. Nalm6^FasKO^ and SupT1^FasKO^ parental cell lines were used for the restimulation experiments.

### Detection of cytokines

Cytokine concentrations in cell culture supernatants were measured by ELISA using kits to detect IFNγ (Biolegend; 430104), IL-2 (Biolegend; 431804), CCL1 (R&D Systems; DY272), CXCL10 (R&D Systems; DY266), CXCL13 (R&D Systems; DY801), according to the manufacturer’s instructions using a Multiskan™ FC microplate photometer (Thermo Scientific).

### Detection of apoptotic cells

Transduced T cells were treated as described, incubated for five hours at 37°C, 5% CO_2_, surface stained for CD3 and RQR8, washed once in PBS, washed once in Annexin V binding buffer (Biolegend; 422201), resuspended in Annexin V binding buffer with Annexin V BV421 (Biolegend; 640924) and incubated for 15 minutes at room temperature protected from the light. Cells were then washed and resuspended in Annexin V binding buffer containing 7-AAD and analysed by flow cytometry. Apoptotic cells were defined as being Annexin V^+^ 7-AAD^−^.

### Immobilised FasL assays

2 μg of recombinant FasL (PeproTech; 310-03H) was immobilised onto 96-well microplates (Starlab; CC7672-7596) overnight at 4°C and the plate was washed several times with PBS. For the proliferation experiments, 5×10^4^ CAR T cells were seeded onto the FasL-immobilised microplate and incubated for five days. Number of CAR T cells were quantified by flow cytometry, using CountBright™ Counting Beads. For the measurement of NF-κB activity, 1×10^5^ transduced NF-κB Jurkat reporter cells were seeded onto the FasL-immobilised microplate, incubated overnight and then treated as described.

### NF-κB reporter assay

1×10^5^ transduced NF-κB Jurkat reporter cells were cultured with immobilised FasL (20 μg/ml) overnight, at which point cells were analysed with the Bright-Glo™ Luciferase Assay System (Promega; E2610) according to the manufacturer’s instructions, and then luminescence measured on a Varioskan™ LUX microplate reader (Thermo Scientific).

### Transcriptomic analysis using the Nanostring platform

2 μg of recombinant FasL (PeproTech; 310-03H) was immobilised onto 96-well microplates (Starlab; CC7672-7596) overnight at 4°C and the plate was washed several times with PBS. 2×10^5^ CAR T cells were then seeded onto the FasL-immobilised microplate and incubated at 37°C, 5% CO_2_ for three days. RNA was extracted from the microplate using the RNAspin Mini Kit (Merck; GE25-0500-71) and quantified using a NanoDrop™ Spectrophotometer (ThermoFisher). 50 ng of extracted RNA was sequenced using the nCounter® CAR-T Characterization Panel (NanoString®) and analysed on the nCounter® SPRINT Profiler (NanoString®) according to the manufacturer’s instructions.

### Memory and exhaustion phenotyping

For memory phenotyping, CAR T cells were stained for CD62L and CD45RA expression, with CD62L^+^CD45RA^+^ being naïve T cells (TN), CD62L^+^CD45RA^−^ being central memory T cells (TCM), CD62L^−^CD45RA^−^ being effector memory T cells (TEM), and CD62L^−^CD45RA^+^ being effector memory T cells expressing CD45RA (TEMRA). For expression of markers associated with exhaustion, CAR T cells were stained for PD-1, LAG3 and TIM3, using Boolean gating to identify cells expressing one, two or three of these markers. To get an accurate representation of the cell’s phenotype, only cohorts that had at least 2000 cells acquired in the CAR T gate (CD3^+^RQR8^+^) on the flow cytometer were analysed. Any cohorts below this threshold were excluded from analysis.

### Immobilised anti-GD2 CAR restimulation assays

100 ng anti-GD2 CAR idiotype antibody (anti-Huk666) was immobilised onto 96-well microplates overnight at 4°C and the plates were washed several times with PBS prior to seeding with 1×10^5^ CAR T cells, which were incubated for three or four days. CAR T-cell numbers were enumerated by flow cytometry using CountBright™ Counting Beads and were reseeded onto another anti-GD2 CAR immobilised microplate.

### *In vivo* studies

All animal studies were performed under a UK Home Office–approved project license. 6–10-week-old female NSG mice (Charles River Laboratory) were raised under pathogen-free conditions. 0.5×10^6^ Nalm6 cells engineered to express firefly luciferase (FLuc) and a HA tag were inoculated intravenously into NSG mice four days prior to CAR T-cell engraftment. Mice were randomized one day prior to CAR T-cell engraftment, where the following day 3×10^6^ CAR T cells were injected intravenously. Tumour engraftment and ongoing tumour growth was measured by bioluminescent imaging (BLI) using the IVIS Spectrum System (PerkinElmer) after intraperitoneal injection of VivoGlo™ luciferin (Promega; P1041). Human T cells were transduced at an MOI of 1.5.

### Data analysis

Data and statistical analyses were performed on GraphPad Prism 9. Flow cytometry analysis was performed on FlowJo (v10.8.1). Transcriptomic analysis from the NanoString platform was performed using nSolver 4.0, R and Python.

## DECLARATIONS

C.M., M.E.K., F.P., M.R., K.L., C.A., J.S., and S.T. are employees and shareholders of Autolus Ltd.

M.P. is a founder of Autolus Ltd, the Chief Scientific Officer, shareholder, and a member of its scientific advisory board.

S.C. is a shareholder of Autolus Ltd.

## AUTHOR CONTRIBUTIONS

**C.M**.: Conceptualization, formal analysis, validation, investigation, visualization, methodology, writing – original draft, writing – review and editing. **M.E.K**.: formal analysis and investigation. **F.P**.: Data analysis and DNA plasmid generation. **M.R**.: Animal model investigation. **K.L**., **C.A**.: DNA plasmid generation. **J.S**.: Cell line generation, writing – review and editing. **S.C**.: Conceptualization, supervision, methodology. **S.T**.: Supervision, methodology, writing – review and editing. **M.P**.: Conceptualization, resources, supervision, funding acquisition, writing – review and editing.

