## Supplementary material for "Novel Fas-TNFR chimeras that prevents Fas ligand-mediated kill and signals synergistically to enhance CAR T-cell efficacy": Figures S1-7

### Figure S1

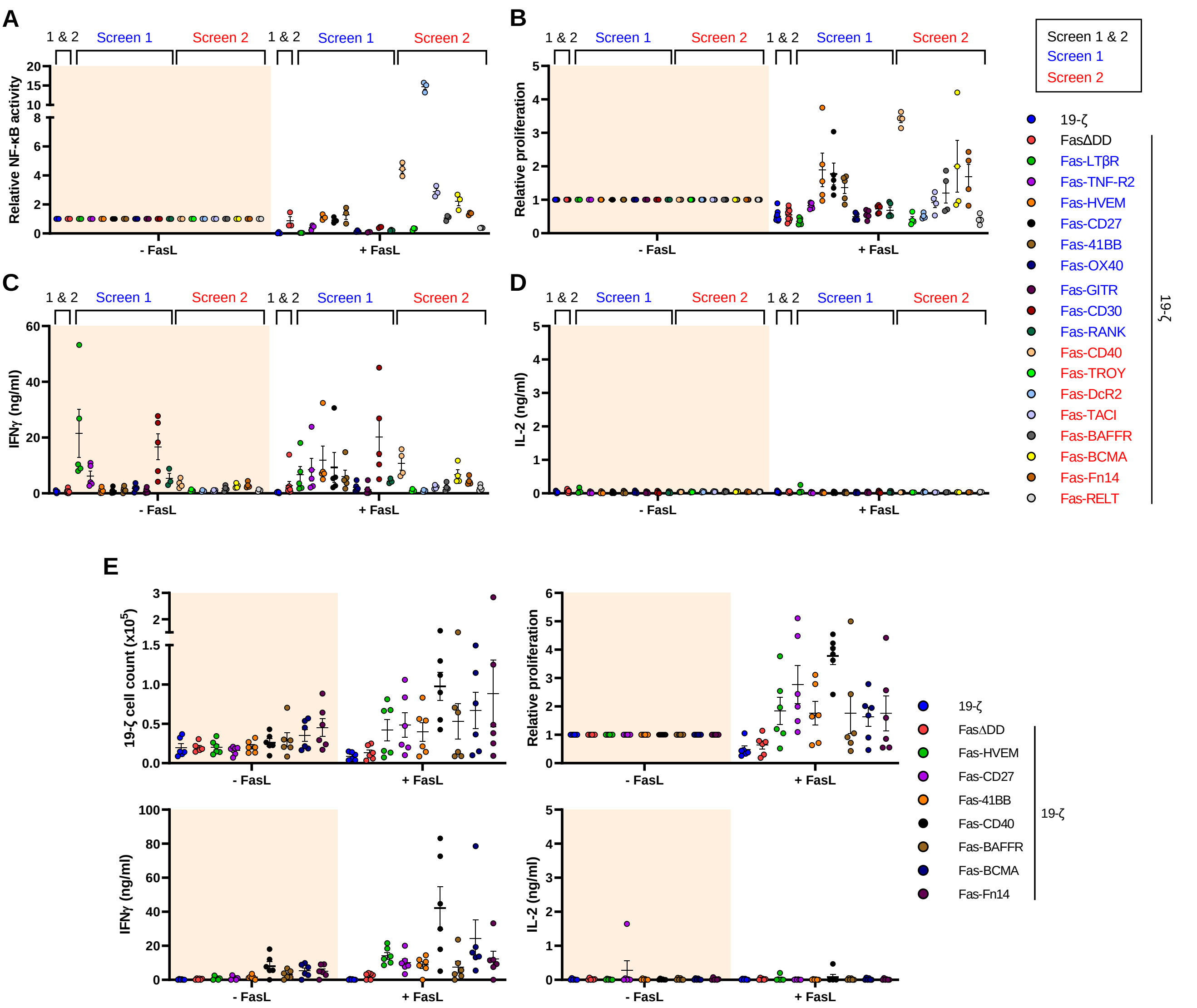


**Figure S1. Fas-TNFRs alter CAR T-cell activity.** (A) Relative NF-κB activity from Figure 1D. Error bars are SEM. (B) Relative T-cell proliferation from Figure 1E. Error bars are SEM. (C and D) After the five-day timepoint from Figure 1E, cell culture supernatant was analysed for IFNγ (C) and IL-2 (D). Error bars are SEM. (E) Fas-TNFR chimeras that induced proliferation upon binding FasL (Figures 1E, S1B) were tested under identical assay conditions using six different independent donors. $5x10^4$ 19-ζ cells were cultured with or without immobilised recombinant FasL (20 µg/ml) for five days, where absolute cell counts (top left) and relative cell counts (top right) were measured, along with IFNγ (bottom left) and IL-2 (bottom right) secretion into the cell culture supernatant. Error bars are SEM.

### Figure S2

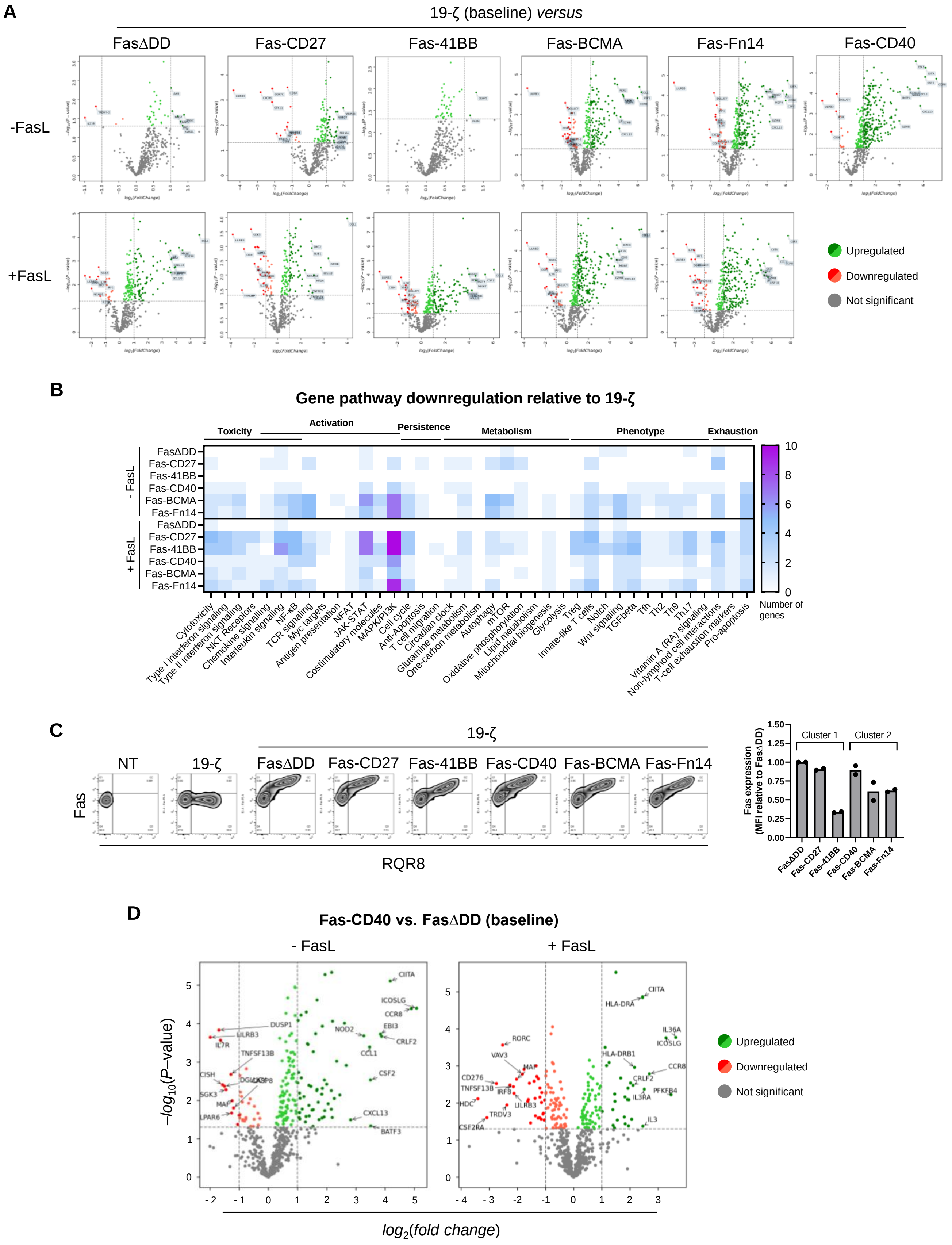


**Figure S2. Fas-TNFRs alter T-cell transcriptome.** (A) Volcano plots of 19-ζ cells co-expressing either FasΔDD or the stated Fas-TNFRs compared to 19-ζ alone, in the presence or absence of immobilised FasL (as described in Figure 1F). (B) Number of significantly ($P < 0.05$) downregulated DEGs relative to 19-ζ were categorised by pathway involvement. (C) Left: Representative flow cytometry plots from one human T-cell donor transduced to express either 19-ζ alone or 19-ζ co-expressing FasΔDD or the stated Fas-TNFRs. Right: median fluorescence intensity (MFI) of the Fas-TNFRs relative to FasΔDD MFI, measured from top right (Q2) quadrant in flow cytometry plots. Two independent donors tested. (D) Volcano plot from experiment described in Figure 1F of Fas-CD40-19-ζ cells compared to FasΔDD-19-ζ cells with or without FasL incubation.

### Figure S3

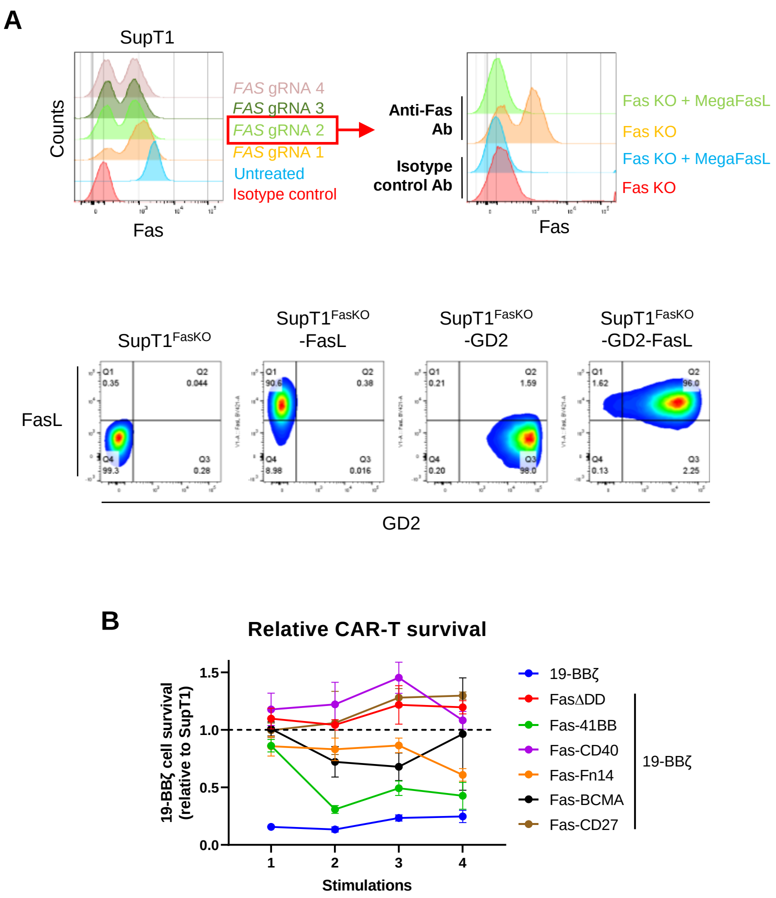


**Figure S3. Generation of SupT1 target cells expressing FasL and GD2.** (A) Top: SupT1 cells were transfected by nucleofection with Cas9 nuclease and four guide RNAs (gRNA) targeting *FAS*, where knockout (KO) efficiency was determined by staining for surface Fas expression. SupT1 cells treated with *FAS* gRNA 2 were then treated with *Mega*FasL (100 ng/ml) to eliminate non-transfected Fas positive cells to create a pure Fas KO population. Bottom: SupT1$^{FasKO}$ cells were transduced to express FasL, GD2 or FasL and GD2. (B) Relative 19-BBζ cell survival after four target cell stimulations from Figure 2E. Error bars are SEM.

### Figure S4

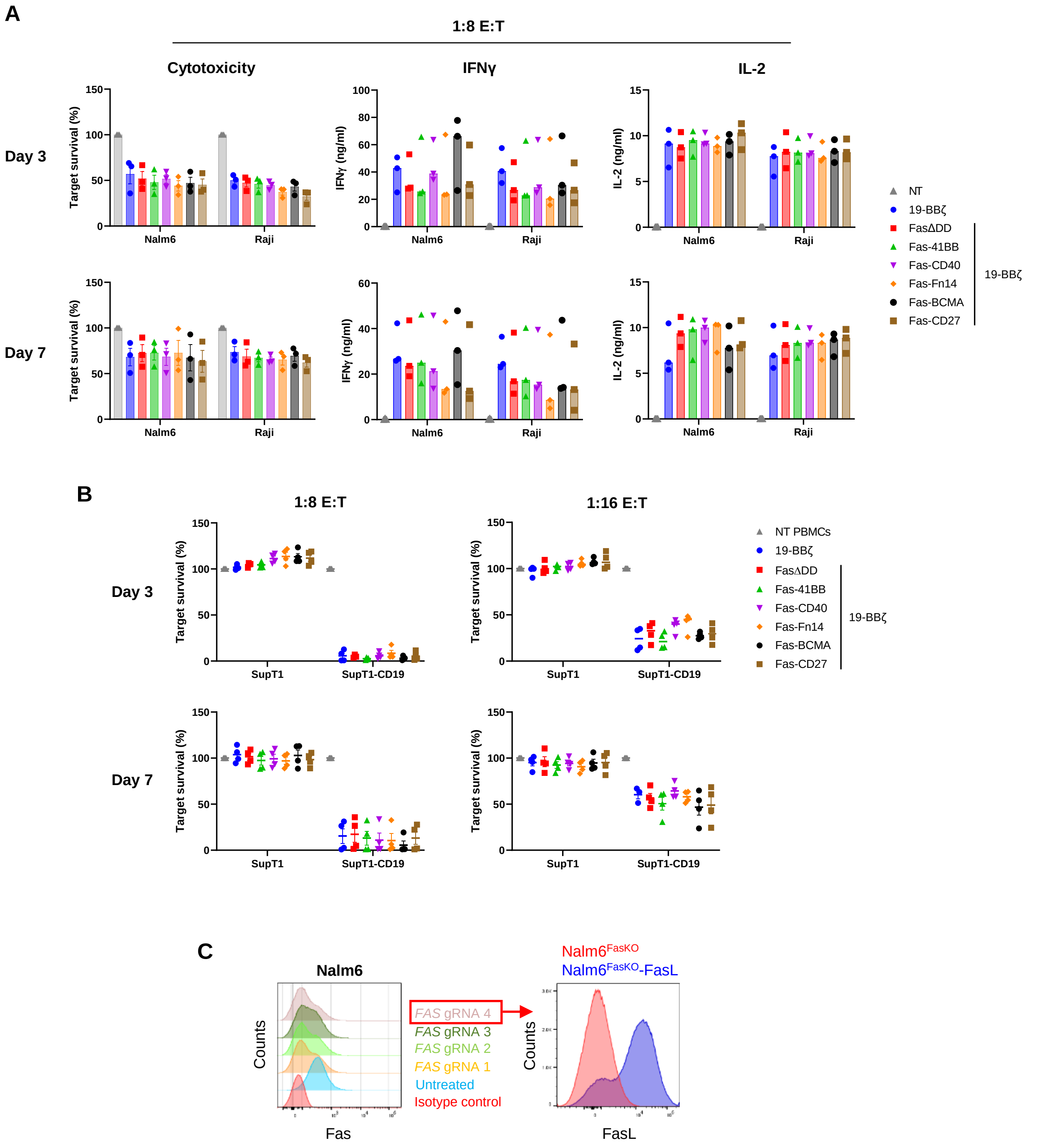


**Figure S4. Fas-TNFRs display equivalent target cytotoxicity after one round of stimulation.** (A) Same experiment as described in Figure 3A, however cells were cultured at a 1:8 E:T. (B) 19-BBζ cells co-expressing FasΔDD or the Fas-TNFRs were cultured with SupT1 or SupT1-CD19 cells for three or seven days, at 1:8 and 1:16 E:Ts, measuring for target survival. Four independent donors tested, error bars are SEM. (C) Left: Nalm6 cells were transfected by nucleofection with Cas9 nuclease and four guide RNAs (gRNA) targeting *FAS*, where KO efficiency was determined by staining for surface Fas expression. Right: Nalm6 cells treated with *FAS* gRNA 4 were then treated with *Mega*FasL (100 ng/ml) to eliminate non-transfected Fas positive cells to create a pure Fas KO population (Nalm6$^{FasKO}$) and were then transduced to express FasL (Nalm6$^{FasKO}$-FasL).

### Figure S5

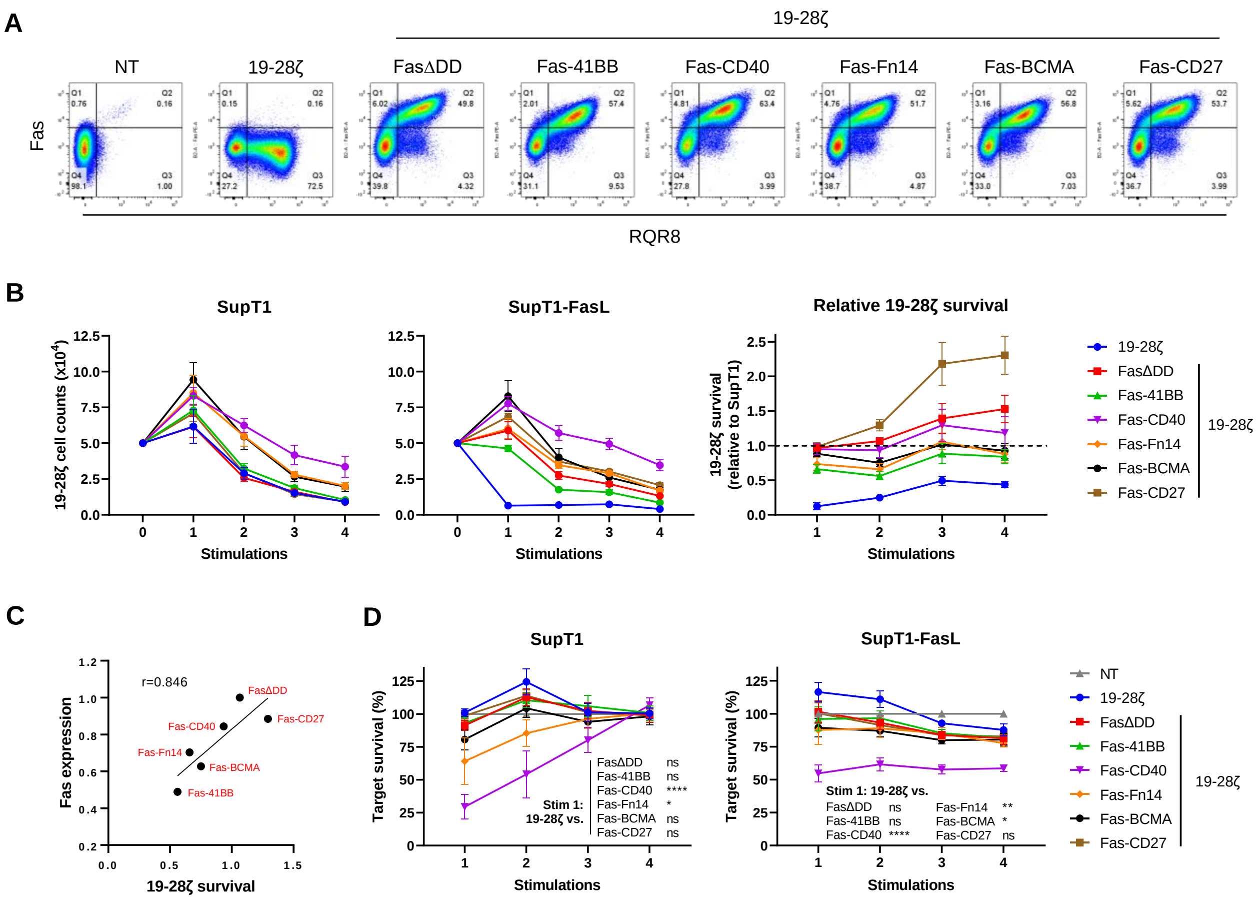


**Figure S5. Fas-CD40 co-expressed with 19-28ζ increases background cytotoxicity.** (A) Representative flow cytometry plots from one human T-cell donor transduced to express either 19-28ζ alone or co-express FasΔDD or the stated Fas-TNFRs. (B) Left and middle: 19-28ζ cells co-expressing FasΔDD or the Fas-TNFRs from four independent donors were stimulated with $5x10^4$ $SupT1^{FasKO}$ or $SupT1^{FasKO}$-FasL cells up to four times, at an initial 1:1 E:T, with cell counts being analysed after each stimulation. Right: Relative 19-28ζ cell survival from the four target stimulations. Error bars are SEM. (C) Mean average of relative Fas expression (from Figure 4B) *versus* mean average of relative 19-28ζ survival (from second stimulation readout in Figure S5B), r = Pearson correlation coefficient. (D) From experiment described in B, the percentage of surviving targets analysed after each target stimulation. Error bars are SEM, $*P < 0.05$, $**P < 0.01$, $****P < 0.0001$, ns – non-significant, two-way ANOVA.

### Figure S6

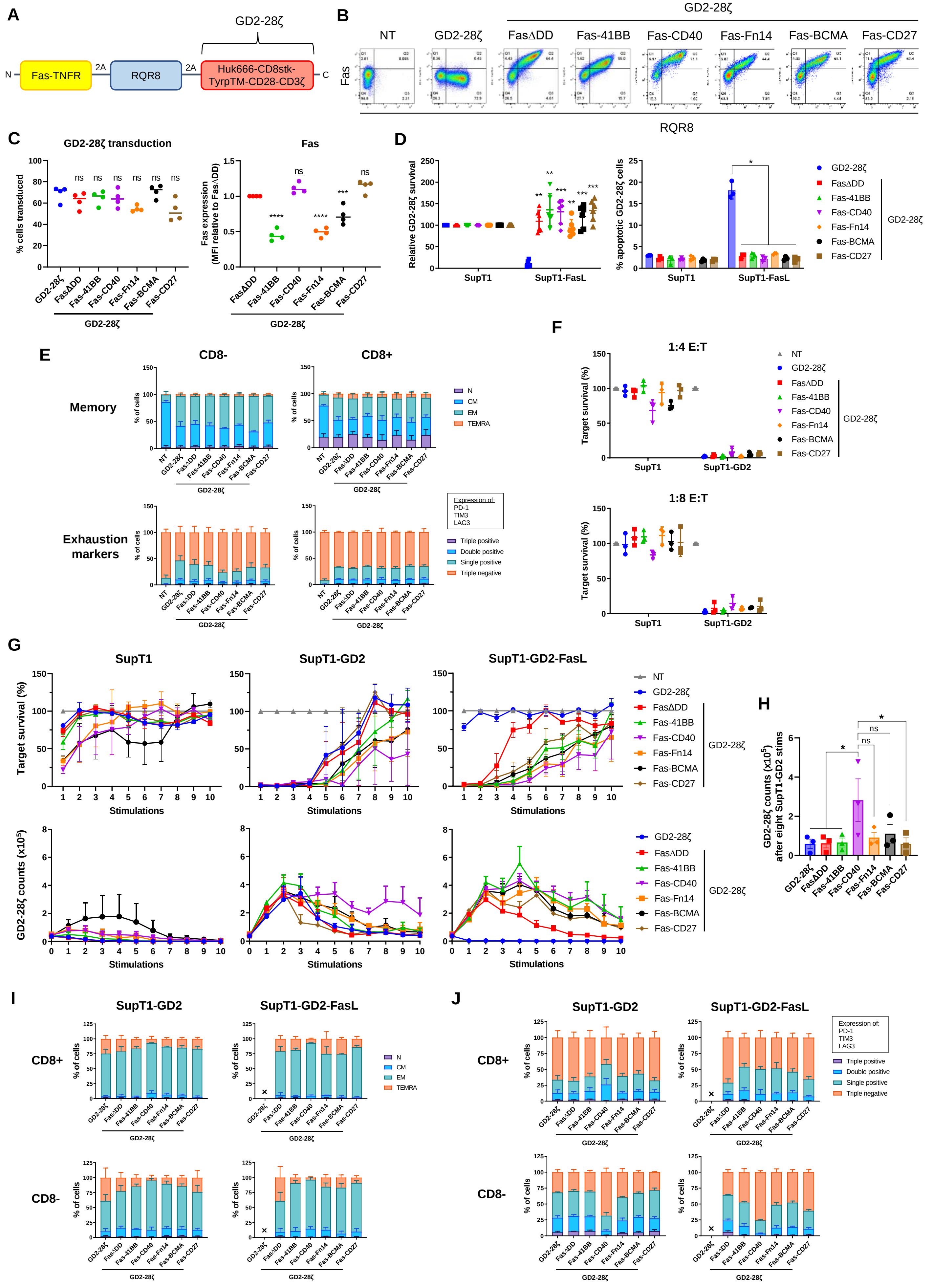
A
GD2-28ζ
N Fas-TNFR 2A RQR8 2A Huk666-CD8stk-TyrpTM-CD28-CD3ζ C
B
GD2-28ζ
NT GD2-28ζ FasΔDD Fas-41BB Fas-CD40 Fas-Fn14 Fas-BCMA Fas-CD27
Fas
RQR8
C
GD2-28ζ transduction
% cells transduced
Fas
Fas expression (MFI relative to FasΔDD)
D
Relative GD2-28ζ survival
% apoptotic GD2-28ζ cells
SupT1
SupT1-FasL
E
CD8-
CD8+
Memory
Exhaustion markers
% of cells
N
CM
EM
TEMRA
Expression of: PD-1 TIM3 LAG3
Triple positive
Double positive
Single positive
Triple negative
F
1:4 E:T
1:8 E:T
Target survival (%)
SupT1-GD2
G
SupT1
SupT1-GD2
SupT1-GD2-FasL
Target survival (%)
GD2-28ζ counts (x10^5)
Stimulations
H
GD2-28ζ counts (x10^5) after eight SupT1-GD2 stims
I
J
CD8+
CD8-

**Figure S6. Fas-TNFRs enhance GD2-28ζ CAR efficacy.** (A) Schematic of polycistronic transgene transduced into human T cells. GD2-28ζ: Huk666 binder fused to the endodomains of CD28 and CD3ζ via a CD8 stalk and Tyrp transmembrane domain. (B) Representative flow cytometry plots from one human T-cell donor transduced to express either GD2-28ζ alone or co-express FasΔDD or the stated Fas-TNFRs. (C) Left: transduction percentages of T cells from four independent donors, ns – non-significant, one-way ANOVA (Dunnett's multiple comparisons test relative to GD2-28ζ). Right: MFI of the Fas-TNFRs relative to FasΔDD MFI, measured from top right (Q2) quadrant in B. Four independent donors tested, mean being shown, ***$P$ < 0.001, ****$P$ < 0.0001, ns – non-significant, one-way ANOVA (Dunnett's multiple comparisons test relative to FasΔDD). (D) GD2-28ζ cells were cultured with SupT1$^{FasKO}$ or SupT1$^{FasKO}$-FasL cells at a 1:1 E:T, either for 72 hours (left) or five hours (right), at which point GD2-28ζ cell survival or percentage of apoptotic cells (Annexin V$^{+}$ 7AAD$^{-}$) were calculated, respectively. Six and three independent donors were tested for the cell survival and apoptotic analysis, respectively. Error bars are SEM, *$P$ < 0.05, **$P$ < 0.01, ***$P$ < 0.001, two-way ANOVA. (E) Memory phenotype analysis (top) and percentage of exhaustion marker expression (bottom; PD-1, TIM3, LAG3) from GD2-28ζ cells co-expressing FasΔDD or the Fas-TNFRs under basal conditions. (F) GD2-28ζ cells co-cultured with SupT1 or SupT1-GD2 (Fas$^{+/+}$) target cells for 72 hours at 1:4 and 1:8 E:Ts, measuring for target survival. Three independent donors tested, error bars are SEM. (G) GD2-28ζ cells from three independent donors were stimulated up to ten times with either SupT1$^{FasKO}$, SupT1$^{FasKO}$-GD2 or SupT1$^{FasKO}$-GD2-FasL cells at a starting 1:1 E:T, measuring for target survival and GD2-28ζ cell counts after each stimulation. Effectors were stimulated with 50,000 targets for all ten stimulations, error bars are SEM. (H) GD2-28ζ cell counts after eighth round of SupT1$^{FasKO}$-GD2 stimulation, as described in G. *$P$ < 0.05, ns – non-significant, two-way ANOVA, error bars are SEM. (I and J) After the tenth stimulation from experiment described in G, T-cell memory phenotype (I) and the percentage of GD2-28ζ cells expressing exhaustion markers: PD-1, TIM3 and LAG3 (J); were analysed. Error bars are SEM, an 'X' denotes where too few cells were present to accurately determine phenotype.

### Figure S7

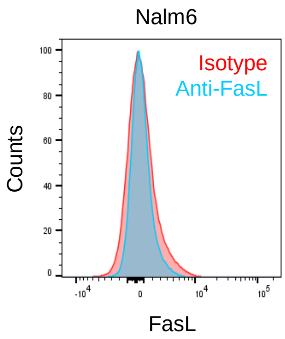


**Figure S7. Nalm6 cells do not express FasL.** Nalm6 cells were surface stained with an anti-FasL antibody or an isotype control antibody.
